# Structural transitions in TCTP tumor protein upon Mcl-1 binding

**DOI:** 10.1101/2022.11.05.515280

**Authors:** Florian Malard, Christina Sizun, Aurélien Thureau, Ludovic Carlier, Ewen Lescop

## Abstract

**Summary:** Translationally Controlled Tumour Protein (TCTP) is a pro-survival factor in tumor cells. TCTP inhibits the mitochondrial apoptosis pathway by potentiating the anti-apoptotic Bcl-2 family members Mcl-1 and Bcl-xL. Specifically, TCTP binds Bcl-xL and inhibits the Bax-dependent Bcl-xL-induced cytochrome c release and TCTP reduces Mcl-1 turnover by inhibiting its ubiquitinylation, thus resulting in decreased Mcl-1 mediated apoptosis. TCTP owns a BH3-like motif forming a β-strand buried in the globular domain of the protein. The crystal structure of TCTP BH3-like peptide in complex with Bcl-xL highlighted the α-helical conformation of TCTP BH3-like motif, suggesting major changes in TCTP structure upon complex formation. However, the structural impact of the interaction on the full-length TCTP and the structural description of TCTP/Mcl-1 interaction are still lacking. Here using biophysical/biochemical methods (NMR, SAXS, circular dichroism, limited proteolysis), we provide an in-depth description of the TCTP/Mcl-1 complex. We demonstrate that full length TCTP binds to the BH3 binding groove of Mcl-1 via its BH3-like motif which interconverts between different binding modes at the micro- to milli-second timescale. As a consequence of the engagement of the BH3-like motif in the interface, the TCTP globular domain is destabilized into a molten-globule state. We also establish that the residue D16 in TCTP BH3-like motif is crucial for the stability and dynamics of the intermolecular interface. As a conclusion, we reveal here in details the structural plasticity of TCTP and discuss its implications for TCTP biology and for future anticancer drug design strategies aiming at targeting TCTP complexes.

**Contact:** Ewen Lescop.

**Supplementary Information:** Supplementary figures, tables and files.

## Introduction

Translationally Controlled Tumour Protein (TCTP) is a small (20 kDa) single-domain globular protein widely conserved across phyla. TCTP, also called fortilin or Histamine Release Factor (HRF), interacts with many partners [1] and is consistently involved in numerous biological processes including tumorigenesis and tumor reversion [2, 3, 4, 5, 6]. TCTP acts as an apoptosis inhibitor and promotes the proliferation of malignant cells. The pro-survival properties of TCTP in cancer cells are supported by numerous evidences such as protection from cell death induced by serum deprivation [7] or cytotoxic drugs [8, 9, 10]. Notably, TCTP is involved in a negative regulatory loop with the major tumor suppressor p53 [11, 12, 13, 14]. TCTP pro-survival properties were also described in the context of interactions with the stress sensor IRE1α [15], the growth factor-beta stimulated clone-22 (TSC) [16] or by inhibition of the Ca^2+^-dependent apoptotic pathways [17].

Importantly, TCTP potentiates central inhibitors of the mitochondrial apoptosis pathway [18], namely Bcl-xL [8, 19] and Mcl-1 [20, 21, 22]. Bcl-xL and Mcl-1 are anti-apoptotic members of the Bcl-2 family proteins and they sequestrate pro-apoptotic factors such as Bax or Bak, respectively, thus inhibiting Mitochondrial Outer Membrane Permeabilization (MOMP) [23]. In mitochondrial-based assays, the molecular interaction between TCTP and Bcl-xL inhibits Bax-dependent MOMP and cytochrome c release. On the other hand, TCTP also exerts its anti-apoptotic role by increasing Mcl-1 stability [20, 22]. Indeed, TCTP inhibits the proteasome-mediated degradation of Mcl-1 by reducing its ubiquitinylation via a yet unknown mechanism [20]. In the context of murine macrophage cell line infection by intramacrophage parasite *Leishmania donovani*, TCTP was found to be strongly associated with Mcl-1, which prevents ubiquitination-mediated degradation of Mcl-1 and inhibits apoptosis [22]. It is well described that Mcl-1 is a target of the E3 ubiquitin-ligase Mule, and that the interaction can be regulated2 by Bcl-2 family proteins which contain a BH3 motif [24]. Hence, TCTP has been proposed to prevent Mule from ubiquinating Mcl-1 to increase the cellular stability of Mcl-1 [22]. Mule also contains a BH3 motif that binds to the BH3 binding groove of Mcl-1 which explains how Bcl-2 family members can regulate Mcl-1 ubiquitination [24, 25]. Therefore, TCTP and Mule might directly compete at the same Mcl-1 surface to dictate Mcl-1 fate. Yet, others have also suggested that Mcl-1 could act as a TCTP chaperone, thus maintaining cellular TCTP levels [21], highlighting the versatile nature of TCTP/Mcl-1 interaction.

TCTP owns a Bcl-2 Homology 3 (BH3) BH3-like motif [19] and is therefore among the few related proteins with anti-apoptotic properties [26]. In the crystal structure of the TCTP BH3-like peptide in complex with Bcl-xL, the peptide binds to the BH3 binding groove of Bcl-xL [19]. The TCTP BH3-like peptide adopts an α-helical conformation in the binding pocket of Bcl-xL, similarly to other structures involving BH3 peptides and Bcl-2 proteins. However, while BH3 motifs are usually disordered or α-helical in the context of the free protein [27], the TCTP BH3-like motif is part of a β-sheet in full length TCTP. Therefore, this suggests important changes in the structure and dynamics of full length TCTP upon complex formation with Bcl-xL [19]. Indeed, the BH3-like motif transition from β-sheet to α-helix is expected, yet not characterized up to date. On the other hand, while the direct interaction between TCTP and Mcl-1 is documented on a functional level [21, 28, 20, 22], it remains unexplored from a structural perspective, which could give indications about how TCTP and Mcl-1 promote their relative cellular levels and functions.

Over the last twenty years, TCTP has become an attractive therapeutical target in cancers [2, 3, 4, 5, 6, 29, 30, 15]. The antidepressant sertraline drug went into phase I/II clinical studies against Acute Myeloid Leukemia (AML), alone or in combination with Ara-C [31]. Sertraline treatment allows reducing TCTP levels and restoring p53 levels in tumor cells, and since sertraline was proposed to directly bind to TCTP, TCTP was suggested to be the cellular target of sertraline [11]. Yet, we recently questioned the direct TCTP/sertraline interaction and proposed alternative modes of action of sertraline in tumor cells [32]. Interestingly, AML often depends on high Mcl-1 levels [7, 33, 34]. Moreover, in classical AML therapy selective inhibitors of Mcl-1 are necessary to prevent the loss of Bcl-xL and subsequent apoptosis in healthy myeloid cells [35]. Therefore, it is important to describe the TCTP/Bcl-xL and TCTP/Mcl-1 complexes from a structural perspective, including the suggested structural reorganization of TCTP, to draw rationals for the design of selective inhibitors.

In this study, we characterized the changes in TCTP structure and dynamics upon complex formation with the Bcl-2 family member Mcl-1 by using a combination of biophysical and biochemical methods (NMR, SAXS, CD, SEC). In addition to revealing details about the major modification in full length TCTP upon complex formation, we used peptidic approaches and molecular docking to highlight the prominent role of the non-canonical residue D16 in the TCTP BH3-like motif for the binding of TCTP to Bcl-2 family partners.

## Material and Methods

### Expression and purification of proteins

#### Production of full length human TCTP

The details of TCTP production and purification were published [36] for ^15^N and/or ^13^C labeled TCTP protein. The protein sequence is given in the supplementary material (Fig. S1).

#### Production of human Mcl-1

The plasmid pET15b (Amp^R^) encoding human Mcl-1 (UniProtKB: Q07820) deleted of the its PEST domain (ΔPEST) and of its C-terminal transmembrane segment (ΔTM) in fusion with N-terminal His_6_-MBP tag was used to express a soluble form of Mcl-1 (ΔPESTΔTM), as described in [37]. A Prescission 3cleavage site was present between His_6_-MBP and Mcl-1. The protein sequence after cleavage is given in the supplementary material (Fig. S1). Unlabeled Mcl-1 ΔPESTΔTM was expressed in *Escherichia coli* BL21 Rosetta (DE3)pLysS grown in rich medium 2xYT. Isotopic labeling was done using minimal medium M9 supplemented with ^15^N-labeled NH_4_Cl (1 g.L^-1^) and ^12^C-α-D-glucose (^15^N-labeling) or ^13^C-α-D-Glucose (^15^N-^13^C labeling). Biomass was grown at 37 °C until OD_600_ ∼ 0.6 and induction of protein expression was achieved using 0.25 mM IPTG at 15 °C during 36 hrs. Bacteria were harvested (5000 g, 20 min.) and pellets resuspended in lysis buffer (50 mM Tris pH 8, 500 mM NaCl, 1 M urea, 2 mM DTT) supplemented with 0.3 mg.mL^-1^ lysozyme and EDTA-free protease inhibitor cocktail (Roche) prior to pressure-assisted lysis using a French press system. After 3 cycles at 1500 bars the lysate was finally centrifuged at 12500 g during 30 min. and the supernatant was collected. This was loaded on a histidine affinity chromatography column (5 mL HisTrap FF crude, GE) equilibrated with the loading buffer (50 mM Tris pH 8, 500 mM NaCl, 1 M Urea and 2 mM DTT) and a flow-rate of 2.5 mL.min^-1^ at 4 °C. The loaded column was washed with 40 mM imidazole (10 CV) and the protein was eluted with 150 mM imidazole (5 CV). The elution product was dialyzed against the Prescission digestion buffer (50 mM Tris-HCl pH 7.4, 150 mM NaCl, 10 mM EDTA and 1 mM DTT) during 2 hours. Digestion was achieved with Prescission protease (1:500 w/w ratio) at 4 °C overnight in order to remove His_6_-MBP tag. The digestion mix was then dialyzed against the loading buffer for histidine affinity chromatography (50 mM Tris pH 8, 500 mM NaCl, 1 M urea, 2 mM DTT). Digested Mcl-1 ΔPESTΔTM was isolated in the flowthrough fraction upon loading the digestion mix onto a histidine affinity chromatography column and washing with the loading buffer (5 CV). The protein was concentrated up to 500 µM and further purified using Superdex 200 10/300 column equilibrated with the storing buffer (20 mM Tris pH 7, 300 mM NaCl, 0.5 mM EDTA, 2 mM DTT). Finally, Mcl-1 ΔPESTΔTM was concentrated up to 500 µM and stored at -80 °C.

#### BH3 peptides derived from TCTP and Mule

Three BH3 peptides derived from TCTP and from the E3 ubiquitine ligase Mule (UniProtKB - Q7Z6Z7) were purchased (Proteogenix) (Fig. S2 A). We designed the peptide TCTP_BH3_ to encompass the BH3-like sequence of TCTP from residue D11 to E32, as defined in [19]. We also designed the TCTP_BH3D16I_ peptide containing the single amino-acid mutant D16I in the BH3 region to restore the hydrophobic residue found in canonical BH3 motifs at this position h1. Finally, we designed the Mule_BH3_ peptide to contain the Mule BH3 region and to have the same length as the above-mentioned TCTP-derived peptides. All peptides were dialyzed against milliQ water in 1 kDa MWCO membrane and vacuum-dried before use in NMR studies.

#### Nuclear Magnetic Resonance

NMR measurements were performed using Bruker AVIII HD 950 MHz and AVIII 800 MHz spectrometers equipped with TCI cryoprobes. For each experiment in this study, detailed experimental conditions are given in the caption of the corresponding figures. Experiments were carried out at 35 °C under acidic conditions (pH 6.5) for peptide studies, while experiments involving full length TCTP and Mcl-1 were performed under alkaline conditions (pH 8) to limit solubility issues upon complex formation and to increase the stability of the complexes.

2D ^1^H-^15^N correlation spectra were collected using the ^15^N SOFAST-HMQC experiment [38]. We completed sequence-specific backbone assignment of ^15^N-^13^C labeled proteins using classical 3D triple resonance experiments with BEST-TROSY principle [39, 40, 41] as implemented in NMRlib pack-_*λ*_ age [42] and on the basis of available assignment data for TCTP [43, 36] and Mcl-1 [37]. The same 3D NMR experiments were used for *de novo* backbone assignment of labeled TCTP and Mcl-1 in complex with partners. 2D and 3D correlation spectra were processed with Topspin 3.5 (Bruker) and analyzed with CCPNMR software 2.4 [44]. Combined ^1^H-^15^N chemical shift perturbations (Δδ_comb_) were calculated according to the equation Δδ_comb_ = (Δδ^1^H + 0.14Δδ^15^N)^1/2^, where Δδ^1^H and Δδ^15^N are the chemical shift perturbations (in ppm) for ^1^H and ^15^N resonances, respectively [45]. For the determination of dissociation constant (K_D_) of the complex of TCTP BH3-like peptide, we used the TITAN line shape analysis software (v1.6) [46] to analyze a titration series of 11 2D ^1^H-^15^N SOFAST-HMQC experiments. Given a fixed ^15^N labeled Mcl-1 concentration (100 µM), the peptide titration points were as follows (µM): 0, 10, 20, 30, 40, 50, 100, 150, 200, 300, 500.

We used a two-state binding model and we defined regions of interest for the following Mcl-1 residues in the TITAN interface: S183, T191, E225, T226, A227, V299, T301. For all selected residues, the corresponding crosspeaks were isolated and exhibited significant chemical shift perturbations along the titration.

#### SEC-SAXS experiments

SEC-SAXS experiments were performed on the SWING beamline at the SOLEIL synchrotron (Saint-Aubin, France). All experiments were done at 37 °C in the following buffer: 50 mM CHES pH 9, 50 mM NaCl and 2 mM TCEP (Table S1). Isolated TCTP and Mcl-1 were prepared at a final concentration of 500 µM. The TCTP/Mcl-1 complex was prepared as an equimolar mix of the two proteins (500 µM each) and incubated for 2 hours. For each sample, a volume of 75 µL was injected onto a size exclusion column (Superdex 200-10/300, GE Healthcare) and eluted directly into the SAXS flow-through capillary cell with a flow rate of 0.5 ml.min^-1^. The SAXS data were recorded using an EigerX 4M detector at a distance of 2 meters with a definition of the momentum transfer 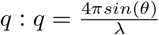 with 2*θ* as the scattering angle, and *λ* the X-ray wavelength (1.033 Å for these experiments). The overall SEC-SAXS setup has already been described previously [47, 48, 49].

In total, 900 SAXS frames were collected continuously upon elution with a frame duration of 1.99 s and a dead time between frames of 0.01 s. 180 frames accounting for buffer scattering were collected before the void volume. Data reduction to absolute units, buffer subtraction and averaging of identical frames corresponding to the elution peak were performed with the SWING in-house software FOXTROT [47] and US-SOMO [50]. US-SOMO was also used to estimate the MW based on the volume of correlation [51]. Data analysis yielding the intensity at *q* = 0(*I*(0)), radius of gyration (Rg) and maximal diameter of the protein (Dmax) were conducted with the PRIMUS software from the ATSAS Suite [52]. Dammif via Primus [53] was also used to compute molecular envelope of free Mcl-1 and in complex with TCTP.

#### Circular Dichroism spectroscopy

The far-UV (180–260 nm) Circular Dichroism (CD) experiments were recorded at 25 °C under N_2_ atmosphere using a JASCO J-810 spectropolarimeter to observe secondary structure content of proteins [54]. The CD cell path length was 0.1 mm. Ten accumulations at 100 nm.min^-1^ with data pitch 0.2 nm and digital integration time (DIT) of 1 s were recorded for each sample. Detector voltage never exceeded 700 V. For simulation of CD spectra, the BeStSel online software (http://bestsel.elte.hu) was used [55]. For each CD experiment in this study, the detail of buffer conditions is given in the caption of the corresponding figures.

#### Protein-peptide docking with HAD-DOCK

We performed molecular docking using HADDOCK 2.2 algorithm [56, 57] to inform on the possible interfaces between Mcl-1 protein and BH3 peptides derived from TCTP. HADDOCK can integrate experimental data such as chemical shift perturbation to predict an accurate model of a protein complex. Therefore, “active” residues were defined by comparing ^15^N SOFAST-HMQC spectra of ^15^N-labeled Mcl-1 in complex with Mule_BH3_ or TCTP_BH3D16I_. Since the sequences of Mule_BH3_ and TCTP_BH3D16I_ are different, it is expected that spectral differences inform on Mcl-1 residues in direct contact with the BH3 peptide ligand (Fig. S3). Correspondingly, the list of Mcl-1 “active” residues provided to HADDOCK interface included: Q221, H252, F254, S255, T259, W261, G262, R263, I264, V265, T266. For BH3 peptides, the list of “active” residues included the positions h1, h2, h3 and h4 (Fig. S2 A). For Mcl-1 and BH3 peptides, the definition of “passive” residues was set to “automatic”. For docking simulation, we used the bound Mcl-1 conformation extracted from the crystal structure of Mcl-1 in complex with Mule BH3 motif [25] (PDB code: 5C6H). For TCTP_BH3_ and TCTP_BH3D16I_ peptides, the α-helical Mule_BH3_ structure was used as a template for further *in silico* mutagenesis in PyMol [58]. Of note, we used an α-helical model for TCTP_BH3_ because it is the conformation found in the crystallographic structure of TCTP_BH3_ in complex with the Mcl-1 homolog Bcl-xL [19].

#### Limited proteolysis

Full length TCTP and its complex with Mcl-1 were analyzed by limited proteolysis in order to evaluate to which extent the proteins remain packed and protected from proteolysis upon complex formation. Experiments were carried out on concentrated samples (500 µM each protein) considering both isolated TCTP or Mcl-1 proteins as well as an equimolar mix of the two. Limited proteolysis was achieved at low (1 µg.mL^-1^) concentration of chymotrypsin or trypsin proteases or with intermediate (5 µg.mL^-1^) protease concentration. As a control of the efficiency of the proteases and to obtain the full degradation pattern, complete digestion was achieved at high (100 µg.mL^-1^) protease concentration. Samples were prepared in 50 mM EPPS pH 8, 50 mM NaCl, 2 mM TCEP and incubated for 6 hours at 37 °C prior to heat-denaturation and SDS-PAGE analysis.

## Results

### The TCTP_BH3_ peptide binds and reorients in the BH3-binding groove of Mcl-1

As introduced before, no structural description of the molecular interaction between TCTP and Mcl-1 is available. Here, we first investigated by NMR spectroscopy if and where TCTP BH3-like peptide (TCTP_BH3_) could bind Mcl-1. NMR and circular dichroism analysis (Fig. S2 B, C) both revealed that TCTP_BH3_ was essentially unfolded in the absence of Mcl-1. We then recorded ^15^N SOFAST-HMQC spectra of ^15^N-labeled Mcl-1 isolated and upon addition of unlabeled TCTP_BH3_ up to 5:1 peptide/Mcl-1 molar ratio (Fig. 1 A). Under these conditions, the formation of Mcl-1/TCTP_BH3_ complex was very fast (*<* min.) and clearly visible through the shift of many Mcl-1 NMR crosspeaks predominantly in the intermediate exchange regime (Fig. S4 A). Using 3D triple resonance experiments, we could assign 96.2 % (152/158 residues) and 86.1 % (136/158 residues) of free and bound Mcl-1, respectively (File S1). The Gly-Pro sequence at the N-terminus, resulting from the TEV cleavage site, plus the proline residues P198 and P289 were not assigned due to the lack of H-N signals. Additional unassigned residues for the apo protein were G219 and F254 in helix α_2_ and the loop connecting α_3_,α_4_, respectively. For the complex, unassigned residues were G219, V220, Q221, F228, Q229, M231 in helix α_2_; M250, H252, F254 in helix α_3_; V258 in the loop connecting α_3_,α_4_; T260, W261, G262, R263, I264, V265 in helix α_4_; L298, F315 in helix α_5_. The increased number of unassigned residues in the complex is due to missing signals in the ^15^N SOFAST-HMQC spectrum, revealing changes in local dynamics. To further map the binding site of TCTP_BH3_, we computed the ^1^H-^15^N combined chemical shift perturbations between free and bound Mcl-1 (Fig. 1 B). We colored the residues on the structure of Mcl-1 accordingly and we highlighted those with dramatic signal extinction in blue (Fig. 1 C). The largest perturbations, including chemical shifts and signal extinction, were consistently clustered within α-helical regions defining the BH3-binding groove of Mcl-1: helices α_2_, α_3_, α_4_ and in the in-between loop α_4_ α_5_ (Fig. 1 C). Of note, we found perturbations in helix α_5_ for residues L298 and T301, not located within the BH3-binding groove of Mcl-1. This is consistent with other reports about Mcl-1 and BH3 motif interactions [59, 60, 61], and it might indicate a conformational change in Mcl-1 upon complex formation with peptides. We also show the location of K194/K197 that are reported as Mule ubiquitination sites (Fig. 1 C) [62]. Those lysines are not part of the BH3 binding region, suggesting that the TCTP_BH3_ binding interface and Mule ubiquitination sites do not overlap. Finally, we used the 2D line shape analysis software TITAN [46] to estimate a dissociation constant (K_D_) of 7.9 ± 0.5 µM for the TCTP_BH3_/Mcl-1 complex (see Material and Methods). This value is consistent with the K_D_ of 12 µM reported by others using Isothermal Titration Calorimetry (ITC) on TCTP_BH3_ complex with Mcl-1 homolog Bcl-xL [19]. Overall, we demonstrated that TCTP_BH3_ peptide binds to the BH3-binding groove of Mcl-1. The peptide likely adopts multiple conformations that exchange at intermediate to slow timescales (µs-ms), leading to exchange broadening of NMR signals from Mcl-1 residues located at the interface in the complex.

**Figure 1:**
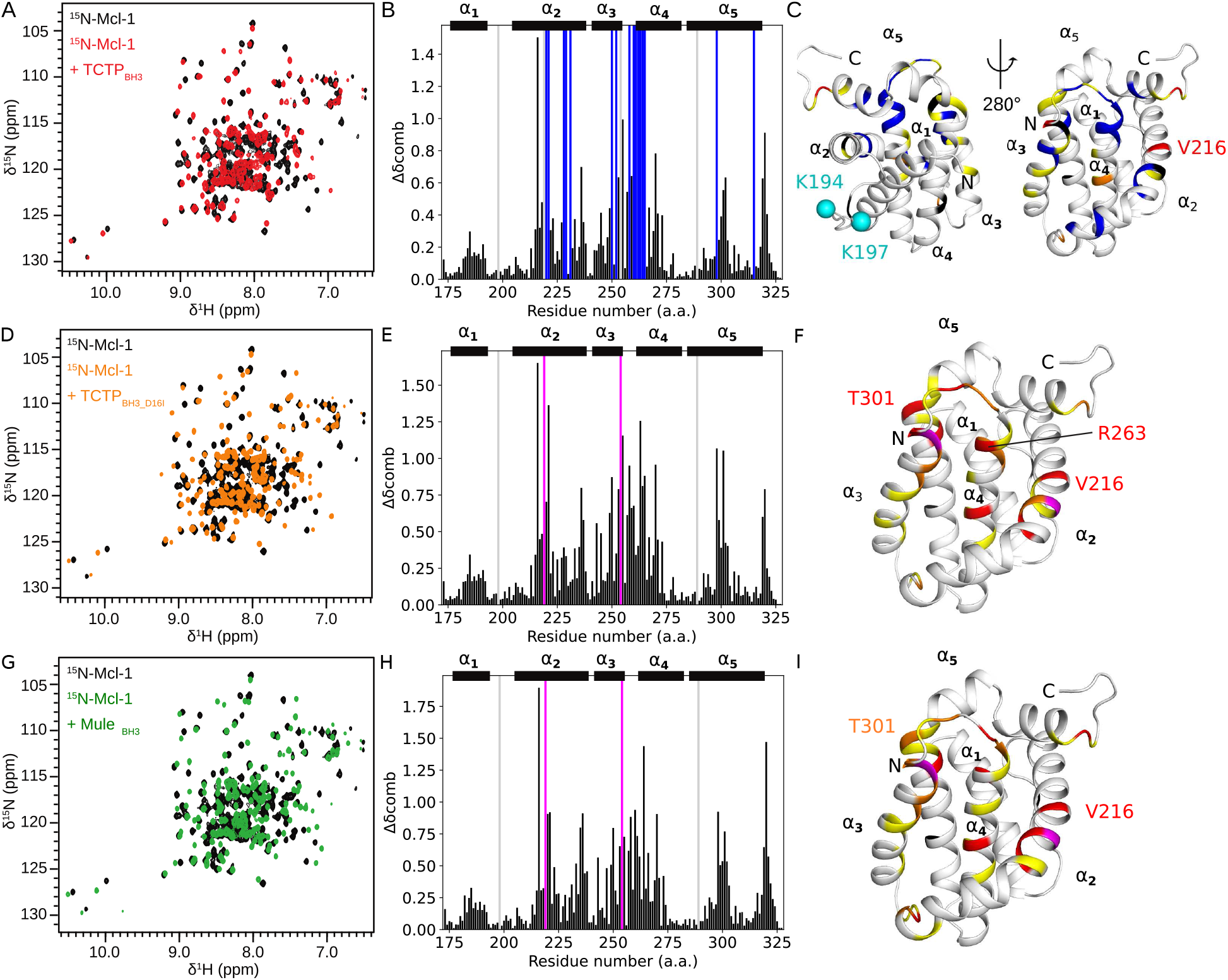
NMR characterization of TCTP_BH3_ and TCTP_BH3D16I_ binding to Mcl-1. (A, D, G) Overlay of ^15^N SOFAST HMQC spectra from isolated ^15^N-Mcl-1 ΔPESTΔTM (Mcl-1) (100 µM, black) and in the presence of excess of unlabeled (A) TCTP BH3-like peptide (TCTP_BH3_) or (D) TCTP BH3-like peptide D16I mutant (TCTP_BH3D16I_) or (G) Mule BH3 peptide (Mule_BH3_). (B, E, H) Combined ^1^H-^15^N chemical shift perturbations in the presence of (B) TCTP_BH3_ or (E) TCTP_BH3D16I_ or (H) Mule_BH3_. Exchange-broadened residues (blue), appearing residues (magenta) and unassigned residues in both free and bound states (gray) are highlighted. (C, F, I) Mapping of combined ^1^H-^15^N chemical shift perturbations in the presence of (C) TCTP_BH3_ or (E) TCTP_BH3D16I_ or (H) Mule_BH3_ (Δδ_comb_ *>* 0.9 ppm, red; 0.9 ppm *>* Δδ_comb_ *>* 0.65 ppm, orange; 0.65 ppm *>* Δδ_comb_ *>* 0.4 ppm, yellow; Δδ_comb_ *<* 0.4 ppm, white); exchange-broadened residues (blue); appearing residues (magenta), and unassigned residues in both free and bound states (black) are highlighted on the NMR structure of Mcl-1 [80]. Reported ubiquitination sites K194 and K197, located away from the BH3-binding groove, are highlighted (cyan) [62]. Experiments were recorded at 950 MHz and 308 K in the following buffer: 50 mM MES pH 6.5, 50 mM EPPS, 50 mM NaCl, 2 mM TCEP in 95 % H_2_O/ 5 % D_2_O.

### TCTP D16 residue controls the binding mode of TCTP BH3 motif to Mcl-1

The BH3-like motif in TCTP has uncommon properties because it exerts anti-apoptotic functions [19]. In contrast to canonical BH3 motifs, it contains a conserved aspartate residue (D16) [1] at so-called position h1 while a hydrophobic residue is generally found at this position. Canonical BH3 peptide usually adopts helical structure in complex with Bcl-2 proteins and the residue at this position creates direct contact with the protein partner. In the case of TCTP_BH3_/Bcl-xL structure [19], the side-chain of D16 creates contact with Q211. In order to assess the role of D16 residue in the binding mode of TCTP with Mcl-1, we designed the single-point mutant TCTP_BH3D16I_ peptide where the canonical BH3 pattern was restored by changing aspartate to isoleucine (Fig. S2 A). As for TCTP_BH3_, circular dichroism analysis showed that TCTP_BH3D16I_ was essentially unfolded in absence of Mcl-1 (Fig. S2 B). Then, we recorded ^15^N SOFAST HMQC spectra of isolated ^13^C-^15^N-Mcl-1 and upon addition of up to 3 molar equivalents of unlabeled TCTP_BH3D16I_ peptide (Fig. 1 D). The complex between TCTP_BH3D16I_ and Mcl-1 was fully formed between one and two molar equivalents of peptide. Under these conditions, all residues assigned in the apo Mcl-1 were clearly visible in the spectrum of bound Mcl-1, in sharp contrast with the complex with TCTP_BH3_. Many Mcl-1 NMR crosspeaks during the titration displayed an intermediate exchange regime (Fig. S4 B). Using triple resonance experiments, Mcl-1 backbone residues in complex with TCTP_BH3D16I_ were all assigned excepted prolines (File S1) including the residues in the BH3 binding groove. The sharp signals indicated a dramatic change in dynamics in the binding interface upon D16I mutation, enabling the recovery of all Mcl-1 NMR signals (Fig. 2 A). We then computed the chemical shift perturbations (Fig. 1 E) that we reported on Mcl-1 structure (Fig. 1 F). Peaks of residues within helices α_2_, α_3_ and α_4_ were largely shifted, confirming that the D16I mutant binds to the BH3-binding groove of Mcl-1. Taken together, the TCTP_BH3_ and TCTP_BH3D16I_ peptides have very similar binding interfaces on Mcl-1. Yet, the presence of an aspartate at position h1 triggers extensive µs-ms motions at the interface, that are essentially quenched when an isoleucine is present at position h1.

**Figure 2:**
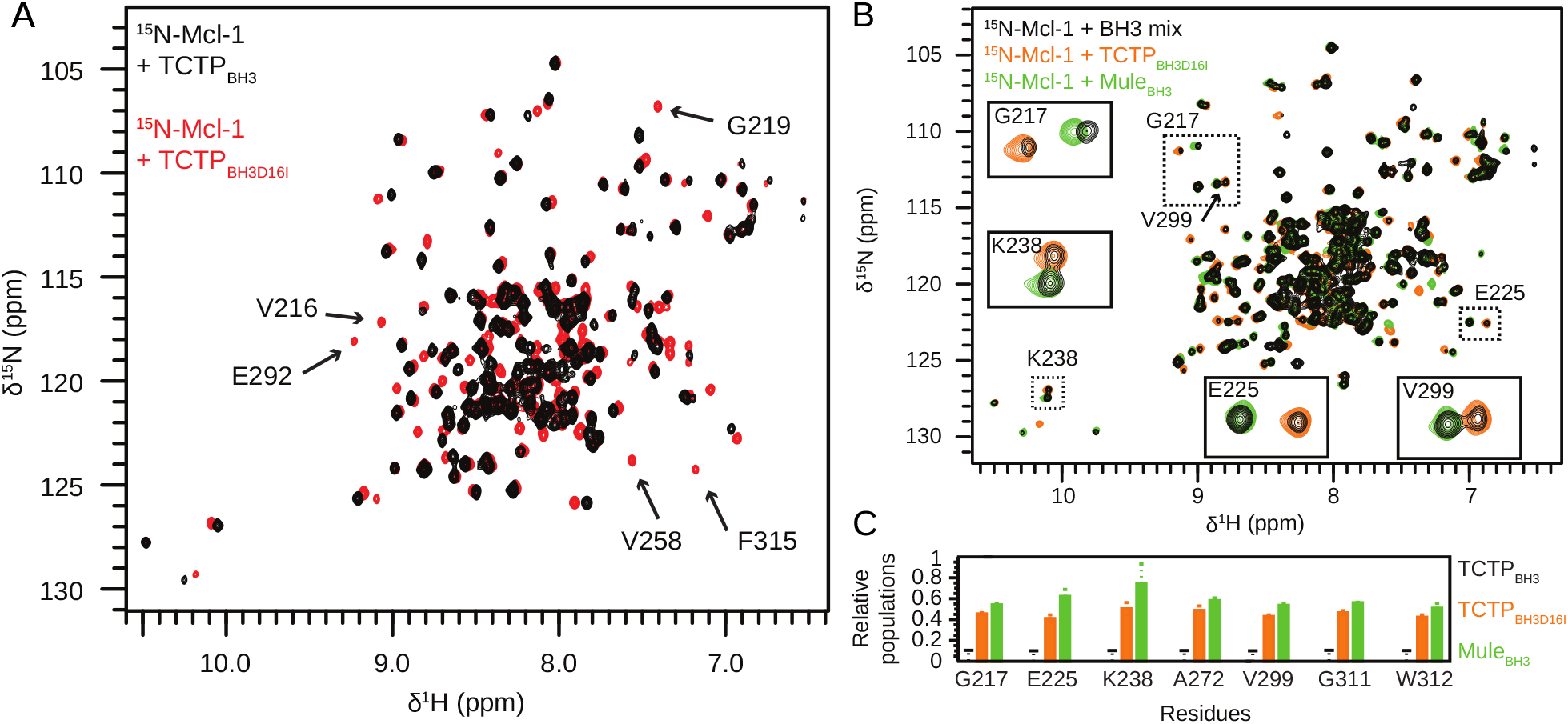
Contribution of TCTP D16 residue in the binding mode to Mcl-1. (A) Overlay of ^15^N SOFAST HMQC spectra from ^15^N-Mcl-1 ΔPESTΔTM (Mcl-1) (100 µM, black) in complex with TCTP_BH3_ (5 eq., black) or TCTP_BH3D16I_ (3 eq., red). (B) Overlay of ^15^N SOFAST HMQC spectra from ^15^N-Mcl-1 in presence of an equimolar mix of BH3 peptides (TCTP_BH3_, TCTP_BH3D16I_ and Mule_BH3_; 5 eq. total, black) or with TCTP_BH3D16I_ (3 eq., orange) or Mule_BH3_ (3 eq., green) alone. (C) Relative population of ^15^N-Mcl-1 bound to TCTP_BH3_, TCTP_BH3D16I_ or Mule_BH3_. Experiments were recorded at 950 MHz and 298 K in the following buffer: 50 mM MES pH 6.5, 50 mM EPPS, 50 mM NaCl, 2 mM TCEP in 5 % D_2_O / 95 % H_2_O.

To compare the binding mode of the TCTP BH3 motif with a canonical BH3 peptide and to gain further insight into the relative roles of TCTP and Mule E3 ligase on Mcl-1, we designed the Mule_BH3_ peptide (Fig. S2 A). As for peptides derived from TCTP BH3-like motif, circular dichroism analysis showed that Mule_BH3_ was essentially unfolded in absence of Mcl-1 (Fig. S2 B). We then carried out an NMR interaction experiment between ^15^N labeled Mcl-1 and up to 3 molar equivalents of Mule_BH3_ (Fig. 1 G). As for TCTP_BH3D16I_, all expected signals, including in the BH3 binding groove, but except prolines, were detectable for Mcl-1 and could be assigned (File S1). We monitored the shift of many Mcl-1 NMR crosspeaks in slow exchange regime during the titration (Fig. S4 C). The complex between Mule_BH3_ and Mcl-1 was fully formed starting from one molar equivalent of peptide, in agreement with the reported dissociation constant (K_D_) of 29 nM for the complex [59]. Then, we confirmed from chemical shift perturbation analysis (Fig. 1 H) that Mule_BH3_ binds to the BH3-binding groove of Mcl-1 (Fig. 1 I). This result is in full agreement with the crystal structure of Mcl-1/Mule_BH3_ complex (PDB code: 5C6H). The strong signals for the BH3 binding groove indicates the absence of significant µs-ms dynamics at Mule_BH3_/Mcl-1 interface, similarly to TCTP_BH3D16I_, in contrast to TCTP_BH3_.

NMR titration experiments revealed a slow exchange regime for Mule_BH3_ peptide, but intermediate regime for TCTP_BH3D16I_ and TCTP_BH3_. Mcl-1 complexes with TCTP_BH3D16I_ or Mule_BH3_ were fully formed at one and two molar equivalents of peptide, respectively. This could indicate stronger affinity of TCTP_BH3D16I_ and Mule_BH3_ peptides than TCTP_BH3_ peptide for Mcl-1. To further explore the thermodynamics of the binding, we performed a competition binding experiment with the three peptides. We prepared an equimolar mix of each of TCTP_BH3_, TCTP_BH3D16I_ and Mule_BH3_ (BH3 mix) and we added an excess of BH3 mix (5 eq.) with ^15^N labeled Mcl-1. We measured ^15^N SOFAST HMQC spectra after 2 hours incubation at room temperature (Fig. 2 B). Based on the known spectra of Mcl-1 bound individually to each peptide (Fig. 1 A, D, G and File S1), we could identify 7 residues as reporter signals that show distinct chemical shifts in the three complexes (G217, E225, K238, A272, V299, G311, W312). All these residues are away from the interface and did not experience significant signal intensity reduction upon peptide binding. For all reporter residues, when the three peptides were mixed, we could detect the spectral signatures from Mcl-1 complex with TCTP _BH3D16I_ and Mule_BH3_ but not for the complex with TCTP_BH3_ (Fig. 2 B, C). This confirmed that TCTP_BH3_ peptide has significantly weaker relative affinity (at least ten times weaker) for Mcl-1 than TCTP_BH3D16I_ and Mule_BH3_. Using the relative signal intensities of the reporter residues as a proxy of the relative populations of the complexes, we demonstrated that the canonical BH3 motifs Mule_BH3_ and TCTP_BH3D16I_ have similar affinity for Mcl-1 with relative population of 55 % and 45 % (Fig. 2 C). These results are fully in agreement with the nanomolar affinity of Mule_BH3_/Mcl-1 complex [59] and the micromolar affinity of TCTP_BH3_/Mcl-1 complex we estimated in the previous section. Overall, the presence of an isoleucine residue at position h1 in TCTP increases the affinity of TCTP for Mcl-1 to a level similar to that of Mule_BH3_. This further indicates that Mcl-1 has a stronger affinity for the BH3 motif of Mule than for that of TCTP. As a conclusion, residue D16 controls both the affinity and the interfacial dynamics of the TCTP/Mcl-1 complex.

### Structural models of Mcl-1 in complex with TCTP_BH3_ or TCTP_BH3D16I_

We next attempted to gain atomic details about the Mcl-1 complex with TCTP BH3-like peptide (TCTP_BH3_). We failed to obtain crystals of the complex. This could possibly be rationalized by the rather low affinity and/or the intrinsic dynamics of the complex. We then resorted to generate structural models from docking experiments using the HADDOCK web interface [63] and the available NMR chemical shift perturbation data. HADDOCK requires submission of a target and ligand, herein structural models of Mcl-1 and BH3 peptide. We submitted the bound Mcl-1 state extracted from the crystal structure (PDB code 5C6H) of the Mcl-1/Mule_BH3_ complex [25]. In three separate runs, we submitted models for α helical Mule_BH3_, TCTP_BH3_ and TCTP_BH3D16I_ peptides. Here, we chose a helical conformation for TCTP-derived peptide, as TCTP BH3 motif adopts an helix in complex with Bcl-xL. Finally, HADDOCK enables molecular docking driven by experimental data. We defined active and passive residues according to our experimental sets of NMR data (see Material and Methods).

For each target-ligand couple, HADDOCK provides a set of clusters and representative structures ranked according to their HADDOCK score, which reflects the relative likelihood of the prediction. In order to validate the docking procedure, we first aligned the cluster ranked first for Mcl-1/Mule_BH3_ to the crystal structure of Mcl-1/Mule_BH3_ complex (pdb code: 5C6H) (Fig. 3 A, S5 A). Overall, the HADDOCK model showed little difference with the experimental structure of Mcl-1 bound to Mule_BH3_ (RMSD Cα = 0.488 Å). Upon alignment of bound Mcl-1 only, the RMSD computed between Mule_BH3_ in the HADDOCK model and the experimental structure was also reasonable (RMSD Cα = 1.186 Å). Moreover, we found a polar interaction between the sidechains of residues D256 (Mcl-1) and Q1981 (Mule) in both HADDOCK model and experimental structure of Mcl-1/Mule_BH3_. Interestingly, the NMR crosspeak for residue D256 in Mcl-1 was very broad in the apo protein, likely because of internal dynamics (File S1). By contrast, the signal from D256 was sharper when Mcl-1 was in complex with Mule_BH3_, suggesting that the internal dynamics of D256 are significantly reduced upon BH3 binding (File S1). The polar interaction of D256 with Q1981 likely stiffens the D256 side chain. Like Mule_BH3_, the TCTP_BH3D16I_ peptide is a canonical BH3 motif because it contains hydrophobic residues at all h1-h4 positions (Fig. S2 A). Therefore, we expect similar binding modes between the top ranked cluster for Mcl-1/TCTP_BH3D16I_ and the crystal structure of Mcl-1/Mule_BH3_ complex (pdb code: 5C6H) (Fig. 3 B, S5 B). Accordingly, we observed that TCTP_BH3D16I_ adopts the canonical binding mode in the BH3 binding groove of Mcl-1 for the top rank cluster, the model showing little difference with Mcl-1/Mule_BH3_. Notably, the side chain of I16 occupies the same hydrophobic cleft in Mcl-1 as V1976 in Mule. As with Mule_BH3_, we observed a sharpening of D256 backbone NMR signal upon complex formation between Mcl-1 and TCTP_BH3D16I_. In the Haddock model, Mcl-1 D256 side chain also forms a salt-bridge with residue R21 of TCTP_BH3D16I_. The side chains of R21 in TCTP and Q1981 in Mule may thus play similar roles by interacting with and stifening D256 dynamics in the respective complexes.

**Figure 3:**
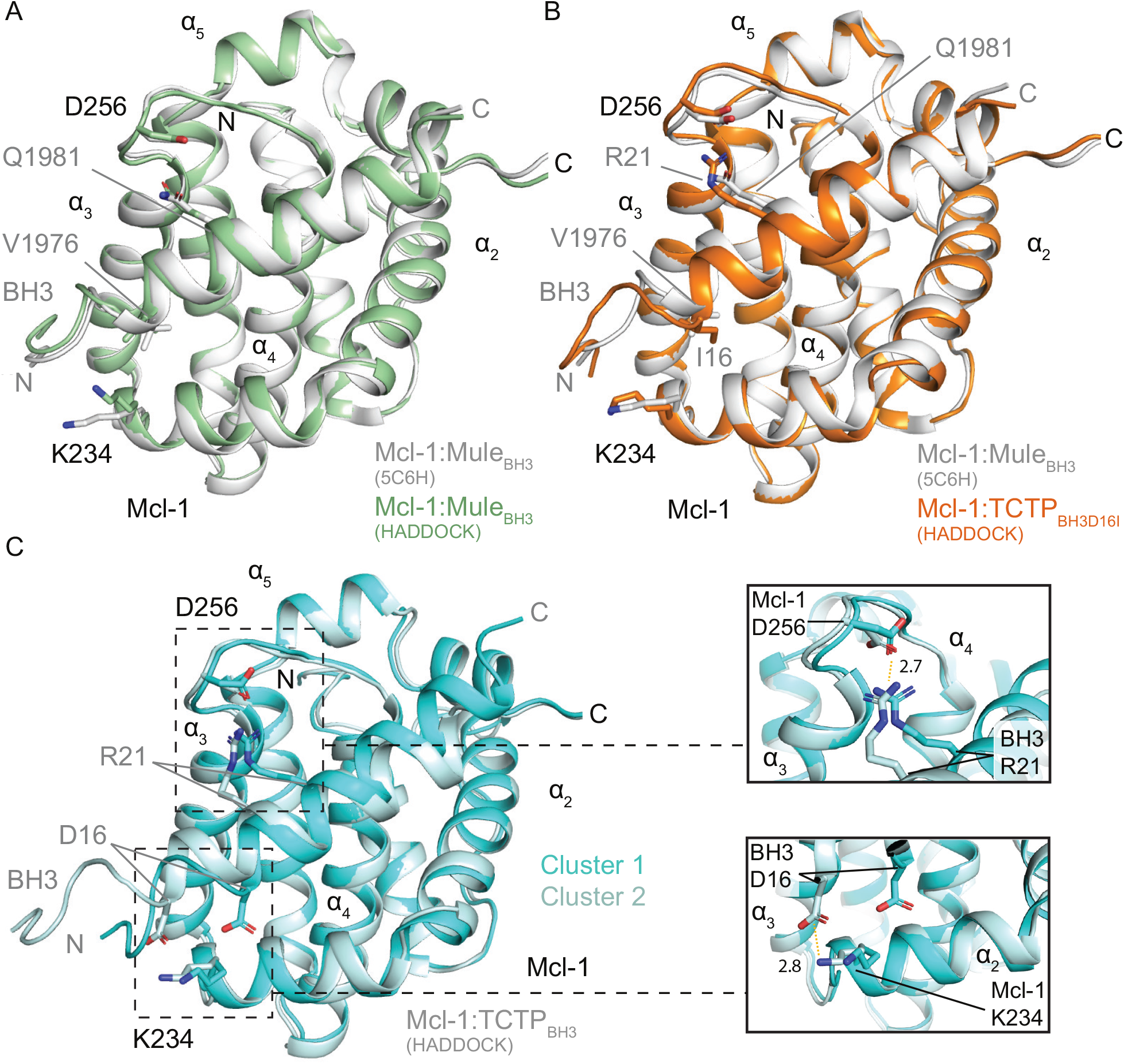
Docking studies of Mcl-1 in complex with BH3 peptides using HADDOCK. (A) Alignment between the representative structure of the top ranked cluster for Mcl-1/Mule_BH3_ and the experimental X-ray structure of Mcl-1/Mule_BH3_ complex (pdb code: 5C6H). (B) Alignment between the representative structure of the top ranked cluster for Mcl-1/TCTP_BH3D16I_ and the experimental X-ray structure of Mcl-1/Mule_BH3_ complex (pdb code: 5C6H). (C) (left) Alignment between the representative structure of clusters 1 and 2 for Mcl-1/TCTP_BH3_. (right) Close-up view of the electrostatic interactions between Mcl-1 D256 with TCTP_BH3_ R21, and Mcl-1 K234 with TCTP_BH3_ D16. Electrostatic interactions are highlighted (dashed orange lines).

In the previous section, we demonstrated that D16 controls both the affinity and the interfacial dynamics of the TCTP/Mcl-1 complex. In order to understand the role of D16 in the BH3-like motif of TCTP, we docked Mcl-1 with an helical peptide containing the TCTP_BH3_ sequence, and we report the alignment of the two best ranked clusters for the complex (Fig. 3 C, S5 C). In the top ranked cluster, the BH3-like peptide adopts a canonical arrangement in the BH3-binding groove of Mcl-1 with D16 occupying a similar position as I16 or V1976 in canonical BH3 peptides. In the second-ranked cluster, the α-helix formed by the BH3-like motif is shifted by 5.6 Å toward the N-terminus of the peptide. In this second-ranked cluster, the h2, h3 and h4 residues occupy the same spatial positions as h1, h2 and h3 in the first-ranked cluster. Despite this shift, the salt-bridge between side-chains of residues R21 in TCTP_BH3_ and D256 in Mcl-1 was preserved in the two clusters. This is consistent with mutagenesis data showing that R21A mutation in TCTP BH3-like motif abrogates binding to Mcl-1 [21]. Notably, we found that in the second cluster the side-chain of Mcl-1 residue K234 forms a salt-bridge with the side-chain of residue D16 in TCTP_BH3_. This suggests that when the protein is in complex with TCTP_BH3_, Mcl-1 K234 side-chain could interact with D16 to create a non-canonical conformational state specific to wild-type TCTP, associated with the global 5.6 Å shift of the BH3 helix within the groove. Hence we propose that the D16 residue in the TCTP BH3-like motif promotes conformational dynamics at the molecular interface by allowing a new non-canonical conformation stabilized by a transient salt-bridge with Mcl-1 K234 in the vicinity of the BH3-binding groove of the protein. The transition between canonical and non-canonical conformations could occur at the µs-ms timescales, as revealed by line-broadening in NMR spectra. NMR line-broadening observed for Mcl-1 residues F228 and M231, located around K234, with TCTP_BH3_, while they are not perturbed upon TCTP_BH3D16I_ binding, also supports this hypothesis (Fig. 1). Conversely, we propose that with the canonical TCTP_BH3D16I_ and Mule_BH3_ sequences, this conformational sampling is quenched, because the hydrophobic residue at position h1 does not allow the formation of the salt bridge with K234.

### Comparison of full-length TCTP and TCTP_BH3_ peptide binding to Mcl-1

In order to compare the binding profiles of full-length TCTP (FL-TCTP) and the truncated TCTP_BH3_ peptide, we next characterized by NMR the interaction between FL-TCTP and Mcl-1. We recorded ^15^N SOFAST-HMQC spectra of isolated ^15^N-labeled Mcl-1 and upon addition of 2:1 excess of unlabeled FL-TCTP (Fig. 4 A). In sharp contrast with Mcl-1/TCTP_BH3_, the formation of the Mcl-1/FL-TCTP complex was slow (∼ hrs) as judged from the slowly appearing cross-peaks corresponding to the complex and disappearing peaks for the apo Mcl-1. In our hands, the stability of the Mcl-1/FL-TCTP complex in solution required higher temperatures and alkaline pH, which is reminiscent of previous observations reported for the TCTP/Bcl-xL complex [19]. In the ^15^N SOFAST-HMQC spectrum of Mcl-1 in complex with FL-TCTP, the visible NMR crosspeaks were generally broadened compared to TCTP_BH3_, in agreement with a complex of larger molecular size (Fig S6 A). As for TCTP_BH3_, several cross-peaks disappeared in the ^15^N SOFAST-HMQC spectrum of Mcl-1 bound to full length TCTP for residues in the BH3 binding groove (Fig 4 B). This strongly suggests that the conformational exchange at intermediate and slower timescales (µs-ms) observed with TCTP_BH3_ is also present in the complex with FL-TCTP. We attempted to assign *de novo* the ^15^N SOFAST-HMQC spectrum of Mcl-1 in complex with FL-TCTP, but we could not obtain exploitable 3D triple resonance spectra.

**Figure 4:**
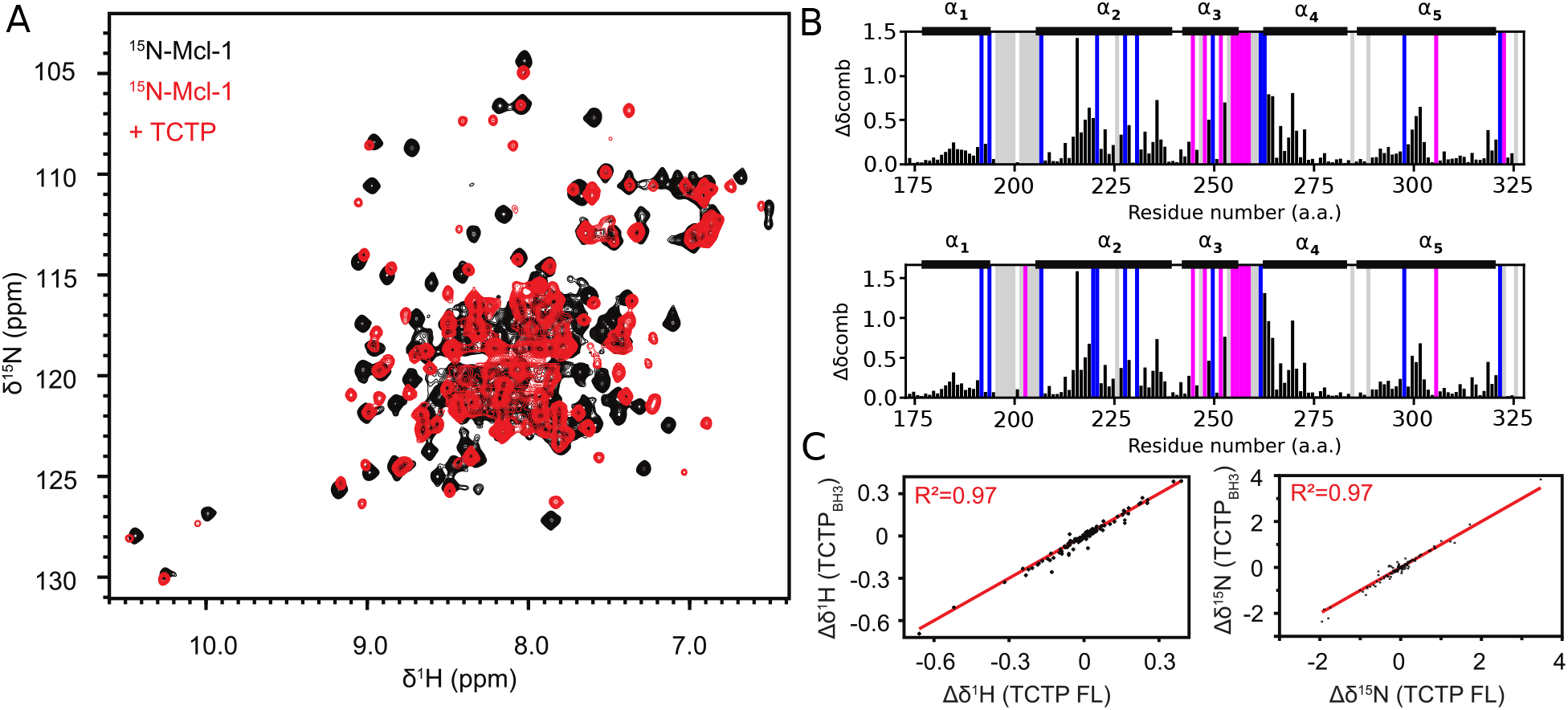
NMR characterization of full-length TCTP binding to Mcl-1. (A) Overlay of ^15^N SOFAST HMQC spectra from isolated ^15^N-Mcl-1 ΔPESTΔTM (Mcl-1) (100 µM, black) and in presence of unlabeled FL-TCTP (2 eq., red) after 2 hours incubation. (B) Alignment of combined ^15^N chemical shift perturbations calculated between isolated ^15^N-Mcl-1 and in complex with (top) TCTP BH3-like peptide or (bottom) FL-TCTP. Largest perturbations were found for residues V216, D218, G219, D236, R263, I264, V265, F270, F273, R300, T301 and K302. Disappearing (K194, R207, V220, Q221, F228, M231, M250, G262 and L298; blue) and appearing (S245, R248, H252, V253, S255, D256, G257, V258 and T259; magenta) residues are highlighted. (C) Correlation of ^1^H (left) and ^15^N (right) chemical shifts perturbations from Mcl-1 in complex with FL-TCTP or TCTP BH3-like peptide. Experiments were recorded at 950 MHz and 308 K in the following buffer: 50 mM EPPS pH 8, 50 mM MES, 50 mM NaCl, 2 mM TCEP in 5 % D_2_O / 95 % H_2_O.

To circumvent these difficulties, we used the same conditions of temperature and alkaline pH required to assemble Mcl-1/FL-TCTP complex to record the ^15^N SOFAST-HMQC spectrum of Mcl-1 in complex with TCTP_BH3_ peptide. The ^15^N SOFAST-HMQC spectra of Mcl-1 bound to TCTP_BH3_ or FL-TCTP were highly resembling in terms of chemical shift profile (Fig S6 B). Therefore, we could easily transfer the chemical shift assignment of Mcl-1 from the complex with TCTP_BH3_ peptide to the complex with FL-TCTP. We show the residue-specific information on ^1^H, ^15^N chemical shift perturbation, signal appearance (magenta) and disappearance (blue) for Mcl-1 in complex with TCTP_BH3_ peptide or TCTP protein (Fig 4 B) and the ^1^H and ^15^N chemical shifts of Mcl-1 in complex with TCTP_BH3_ plotted versus the complex with FL-TCTP (Fig 4 C). Taken together, ^1^H and ^15^N chemical shift analysis reveal that full length TCTP binds the BH3-binding groove of Mcl-1 very similarly to the TCTP_BH3_ peptide, with a similar binding interface and conformational exchange at the molecular interface. Moreover, these results suggest that the interacting region in TCTP is limited to the BH3-like motif and does not involve other regions of TCTP. While sharing a similar binding mode to Mcl-1, TCTP_BH3_ peptide and FL-TCTP protein display a striking difference in their association kinetics with Mcl-1. The slow (∼ hrs) formation of the complex with FL-TCTP strongly suggests a rate-limiting structural reorganization of TCTP prior to complex formation.

To compare the relative affinity of FL-TCTP and TCTP_BH3_ for Mcl-1, we carried out a competition experiment by NMR (Fig. S7). First, the ^15^N labeled FL-TCTP/Mcl-1 complex was prepared, which leads to the disappearance of most signals in ^1^H-^15^N correlation spectra for TCTP. Then, TCTP_BH3_ peptide was added, and most free FL-TCTP signals were recovered (Fig. S7 A). In the reciprocal experiment, the TCTP_BH3_/Mcl-1 complex was prepared, leading to chemical shift perturbation and line broadening of 1D methyl signals for Mcl-1. Then, FL-TCTP was added and no change in spectral signatures from Mcl-1 bound to TCTP_BH3_ could be observed after equilibration time, neither for methyl signatures of free FL-TCTP (Fig. S7 B). This demonstrates that the BH3 peptide of TCTP competes with FL-TCTP at the same binding site on Mcl-1, with stronger affinity, and that the conformational change in TCTP after Mcl-1 binding is reversible. Due to the very slow kinetics of complex formation FL-TCTP, we did not attempt to precisely measure the affinity of the complex.

### Conformational change of TCTP upon binding Mcl-1

In order to characterize the TCTP/Mcl-1 complex from the TCTP perspective, we recorded ^15^N and ^13^C SOFAST-HMQC spectra of isolated ^15^N-^13^C-labeled TCTP and upon complex formation with unlabeled Mcl-1 (Fig. 5 A). In both spectra of isolated TCTP, all crosspeaks were well dispersed in both ^1^H and ^15^N or ^13^C dimensions, indicating that the visible protein is well folded in our conditions. In a previous study, we reported Secondary Structure Propensity (SSP) calculated from Cα, Cβ and Hα chemical shifts using the same TCTP construct and nearly identical experimental conditions [36], well in agreement with the solution structure of full length TCTP [43]. After Mcl-1 was added and equilibrium was reached (2 eq., 2 hours), most cross-peaks from the globular domain of TCTP became invisible due to the complex formation, while strong cross-peaks remained in spectral regions characteristic of disordered residues. Using 3D triple resonances experiments, these visible cross-peaks were easily assigned to the continuous stretch of residues V31 to V70, that encompasses the long internal loop and β-strands β_4_ and β_5_ in the structure of the free protein (Fig. 5 B). We further analyzed ^13^C backbone chemical shifts using the Secondary Structure Propensity (SSP) approach [64] to assess secondary structures changes in the bound TCTP segment visible by NMR (Fig. 5 C). Positive or negative SSP values reveal propensities for α-helical or β-strand conformations, respectively. When comparing SSP for each residue, we observed that values remained unchanged for the loop central region from residue E40 to T65 between free and bound states, suggesting that the loop remains fully disordered in the complex. By contrast, residues in the segments from residues V31 to R38 and residues V66 to V70 underwent an inversion in secondary structure propensity, suggesting a structural transition from β-strand to α-helical conformational propensity. In the free protein, they encompass respectively the strands β_4_ and β_5_, that form a β-sheet at the bottom of the loop. Therefore, this β-sheet breaks upon complex formation with Mcl-1, and residues V31-R38 and V66-V70 become largely flexible, with some helical propensity. Notably, the BH3-like region in TCTP (S15-L29) remained invisible in NMR experiments and therefore the transition from β-sheet to α-helix could not be directly evaluated by NMR. The helical propensity observed for neighboring residues (after V31) may reflect the extension of the putative BH3 helix beyond the BH3 motif when bound in Mcl-1 groove.

**Figure 5:**
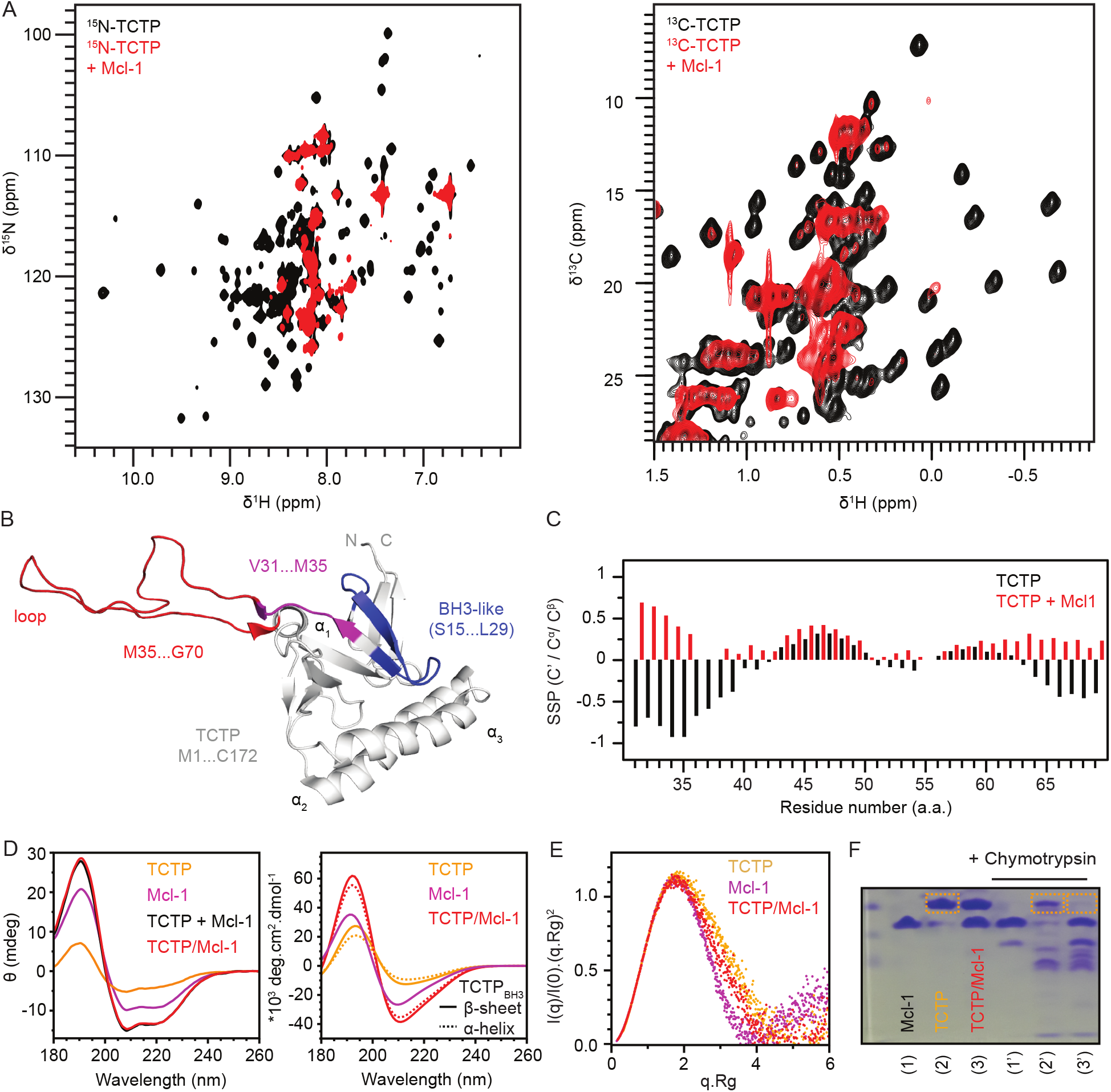
Characterization of molten-globule state of TCTP in complex with Mcl-1. (A) Overlay of (left) ^15^N and (right) ^13^C SOFAST HMQC spectra from isolated ^15^N-TCTP (100 µM, black) and in presence of unlabeled ΔPESTΔTM (Mcl-1) (2 eq., red) after 2 hours incubation. Experiments were recorded at 950 MHz and 278 K in the following buffer: 50 mM EPPS pH 8, 50 mM MES, 50 mM NaCl, 2 mM TCEP in 5 % D_2_O / 95 % H_2_O. (B) The region of TCTP (V31 to G70), for which NMR backbone signals could be assigned, are represented on the NMR structure of the protein [43]. The long flexible loop (red), the BH3-like motif (blue) and its C-terminal extension visible by NMR (magenta) are highlighted. (C) Secondary Structure Propensity (SSP) [64] computed for isolated TCTP (black) and in complex with Mcl-1 (red). Positive values indicate α-helix propensity whereas negative values indicate β-sheet propensity. (D) (left) Far-UV CD experiments on isolated TCTP (50 µM, orange), isolated Mcl-1 (50 µM, magenta), sum of the two CD curves (black) and experimental CD curve of the equimolar mix of the two proteins (50 µM each, red) after 2 hours incubation. Experiments were performed at 308 K in 5 mM phosphate buffer pH 8 and 100 µM TCEP to ensure data reliability at low wavelength. (right) Theoretical prediction of CD curves using BeStSel [55] for native TCTP (orange), TCTP with BH3-like motif as α-helix (dashed, orange), Mcl-1 (magenta). Sum of Mcl-1 and native TCTP curves (red) or with BH3-like motif as α-helix (dashed, red). (E) Normalized Kratky plots for TCTP (orange), Mcl-1 (magenta) and the dominant heterodimeric TCTP/Mcl-1 complex (red). (F) Limited proteolysis experiments2u4sing chymotrypsin protease. Isolated Mcl-1 (500 µM), TCTP (500 µM) and an equimolar mixture were incubated overnight at 308 K in 50 mM EPPS pH 8, 50 mM MES, 50 mM NaCl and 2 mM TCEP with or without chymotrypsin (5 µg.mL^-1^). Dashed line rectangles highlight the band for native TCTP in relevant conditions.

The most striking feature of this NMR experiment is the disappearance of most NMR signals for TCTP residues outside the V31-V70 region. The absence of visible signals for the BH3 motif is likely associated to the extensive µs-ms dynamics at the interface as visible on the Mcl-1 side. However, vanishing signals for residues 1-14 and after 71, i.e. the core protein region, needs to be explained. Severe NMR line-broadening can be expected for large protein complexes or in case of internal dynamics. We therefore decided to complete the NMR studies by complementary structural analysis.

We first relied on far-UV (180-260 nm) Circular Dichroism (CD) spectroscopy to evaluate how secondary structure may be globally impacted in the TCTP/Mcl-1 complex with respect to isolated proteins (Fig. 5 D). The far-UV CD spectrum of isolated TCTP contains contributions from α-helix, β-strand and random coil elements as expected from the crystal and NMR structures of the protein [65, 43]. The far-UV CD spectrum of Mcl-1 only contained the contributions of α-helices, in full agreement with its 3D structure. The experimental CD curve for the complex was compared with the sum of the CD curves for the two isolated proteins TCTP and Mcl-1. The two curves were strikingly similar, indicating that the secondary structure content in TCTP/Mcl-1 complex is closely preserved with regard to isolated proteins. Of note, the putative β-strand to a α-helix transition for the BH3-like motif in TCTP is likely not to be seen herein, since it contributes to less than 5 % of total residues in TCTP/Mcl-1 complex. Overall, CD analysis show limited changes in TCTP and Mcl-1 secondary structure upon complex formation.

### Quaternary structure of the FL-TCTP/Mcl-1 complex

In order to evaluate the stoichiometry of the TCTP/Mcl-1 complex, and hence its molecular size, we performed SEC-SAXS experiments. In SEC experiments, elution time was longer for the isolated Mcl-1 (11.3 min) than for the isolated TCTP (9.8 min), consistent with the molecular weights of 17.9 kDa for Mcl-1 and 19.6 kDa for TCTP. Upon complex formation, a major molecular species with higher apparent molecular weight eluted first (8.5 min.), while residual free TCTP and Mcl-1 could still be observed (Fig. 6 A), revealing incomplete complex formation. We next analyzed SAXS frames corresponding to the major species, presumably TCTP/Mcl-1 complex (Fig. 6 B). We found a molecular weight of 34.7 ± 1.9 kDa and a radius of gyration (R_g_) of 2.22 ± 0.09 nm. We expected a molecular weight of 37.4 kDa for the heterodimeric TCTP/Mcl-1 complex. Considering that the experimental value might be underestimated because residual quantities (*<* 5 %) of free TCTP co-elutes with the complex, the experimental molecular weight of 34.7 ± 1.9 kDa is consistent with a heterodimeric (1:1) TCTP/Mcl-1 complex. In addition, the experimental R_g_ of 2.10 ± 0.08 nm is compatible with a ∼35 kDa globular protein complex. We concluded that the TCTP/Mcl-1 complex is predominantly heterodimeric in solution.

**Figure 6:**
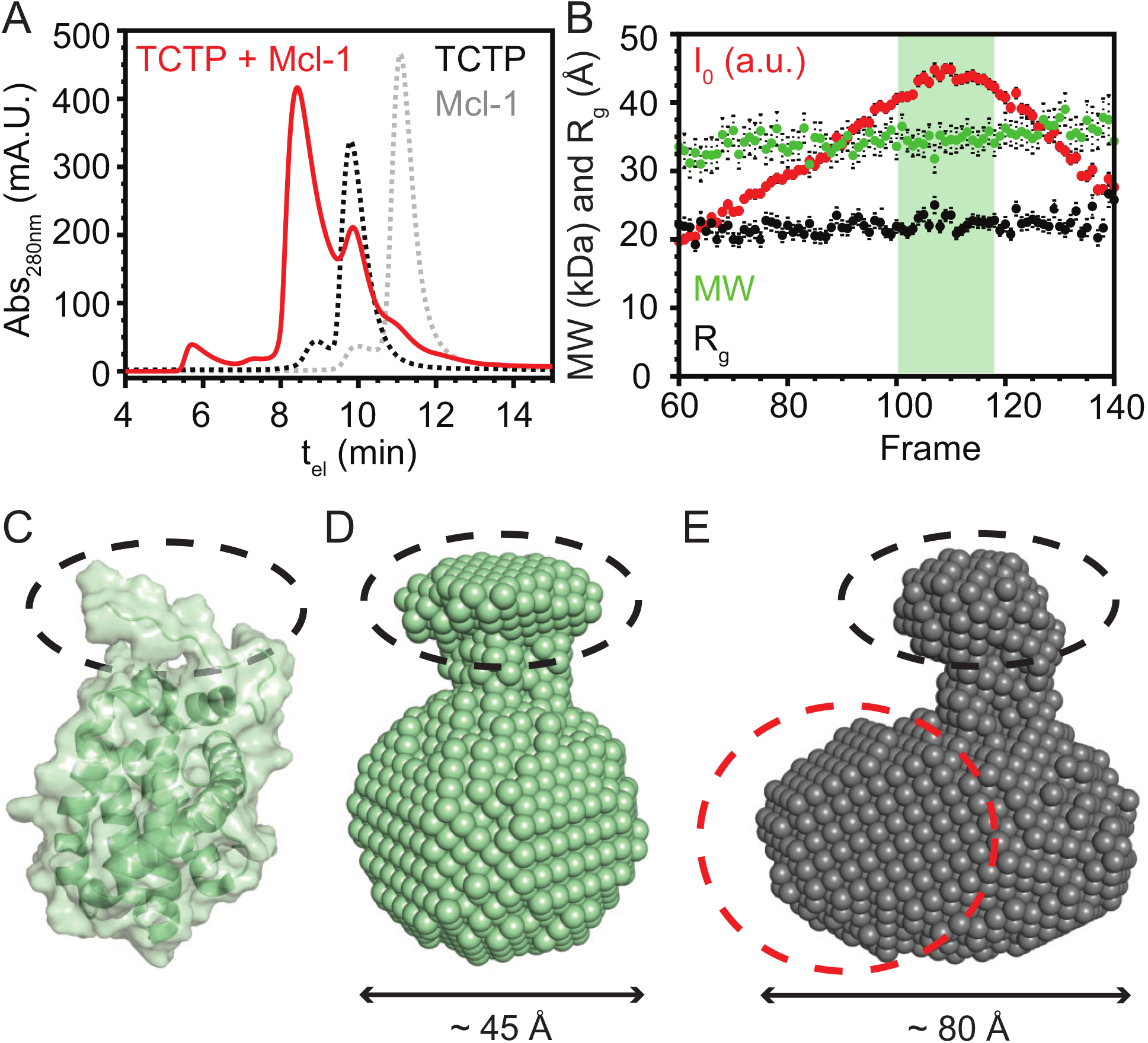
Quaternary structure of TCTP/Mcl-1 complex from SEC-SAXS experiments. (A) SEC profiles of isolated full length TCTP (500 µM, dashed black line), isolated ^15^N-Mcl-1 ΔPESTΔTM (Mcl-1) (500 µM, dashed gray line) and an equimolar mix of TCTP and Mcl-1 after 2 hours incubation (red). (B) SAXS-derived parameters for TCTP/Mcl-1 complex. Scattering intensity I_0_, the molecular weight (MW) and the radius of gyration (R_g_) are shown for the different frames along the SEC dimension. (C) Surface representation of Mcl-1 NMR structure (PDB code 2MHS, [80]). (D) Molecular envelope of free Mcl-1 derived from SAXS analysis. (E) Molecular envelope of free Mcl-1/TCTP complex derived from SAXS analysis. The plausible location of TCTP is highlighted as a dashed red circle. For (C), (D) and (E), the structural signature of Mcl-1 visible as a protuberance pointing out from the globular domain is highlighted as a dashed black ellipse. Experiments were performed at 308 K in the following buffer: 50 mM CHES pH 9, 50 mM NaCl, 2 mM TCEP.

We further exploited SEC-SAXS data by computing molecular envelopes using the Primus software DAMMIF. We identified a potential structural signature for the isolated Mcl-1 that we highlighted on the structure of the protein (Fig. 6 C) and the experimental envelope we computed (Fig. 6 D). Interestingly, we could also detect this signature in the molecular envelope of TCTP/Mcl-1 complex, suggesting a location for Mcl-1 and thus TCTP in the heterodimeric complex (Fig. 6 E). Overall, the TCTP/Mcl-1 complex had a prolate shape with a tightly packed organization, which confirmed the 1:1 binding stoichiometry. More details were not evaluated because residual quantities (*<* 5 %) of free TCTP co-eluted with the complex in SEC, making it difficult to obtain deconvoluted SAXS data with optimal signal-to-noise ratio. Since density was missing for the long internal loop of TCTP in the molecular envelope of the complex, this region may retain its flexible character in the complex.

Molecular envelope computation poorly captures the presence of flexible regions, which are better detectable by the normalized Kratky analysis (Fig. 5 E). We found that the Kratky curve for the isolated Mcl-1 had a typical sine-bell shape that represents a globular, well folded protein. The curve of isolated TCTP [1] represents a globular domain well, but intensity values decreased slower with respect to the diffusion angle q compared to the Mcl-1 curve. This is a signature of an extended segment which can be correlated to the TCTP loop region (not present in Mcl-1). Strikingly, the Kratky plot of TCTP/Mcl-1 complex is highly similar to the mean of curves from isolated proteins, indicating that TCTP retains a fairly packed tertiary structure in complex with Mcl-1 with the exception of the TCTP loop region that is flexible in free and bound states. This corroborates the previous NMR analysis. We concluded that TCTP and Mcl-1 assemble in a tightly packed heterodimeric complex in which both proteins retain properties related to their individual tertiary structure.

### Molten-globule properties of TCTP in complex with Mcl-1

NMR analysis revealed sharp signals for Mcl-1 but extensive broadened signals for TCTP in the complex. This indicated that TCTP and Mcl-1 had significant different dynamic behaviors in the complex. Yet, SEC-SAXS analysis revealed a predominant 1:1 complex in solution. This indicates that line-broadening in TCTP can not be accounted for by formation of large particles. To conciliate all data, we hypothesized that line-broadening in TCTP could originate from the destabilization of the core domain of TCTP into a so-called molten-globular state. Molten-globular states are compact conformational states with tertiary structures that fluctuate at the intermediate-to-slow timescales (µs-ms), while secondary structure elements remain essentially preserved. SEC-SAXS and CD analysis were indeed consistent with a compact nature of TCTP/Mcl-1 with well-formed secondary structures. As a consequence of extensive internal structural fluctuations in the molten-globule state, peptide bonds may be exposed to solvent while they would not in the native state. Therefore, we performed limited proteolysis experiments with chymotrypsin to evaluate if the globular domain of TCTP could be more sensitive to proteolysis upon complex formation (Fig. 5 F). We first established the stability of isolated proteins and complex over time, without chymotrypsin addition, by SDS-PAGE. Then, we obtained full digestion profiles using high chymotrypsin concentration (100 µg.mL^-1^). Finally, limited proteolysis with chymotrypsin was carried out (5 µg.mL^-1^). For isolated Mcl-1, the limited digestion profile was made of a major band corresponding to the undigested protein and a minor band corresponding to one digestion product. For isolated TCTP, a major band corresponding to the undigested protein was also seen along with multiple minor bands for digestion products. For TCTP/Mcl-1 complex, undigested Mcl-1 remained while the band corresponding to undigested TCTP could not be seen anymore. This indicates that peptide bonds are more exposed to solvent in bound TCTP. Overall, we demonstrated that the globular domain of TCTP in complex with Mcl-1 is characteristic of molten-globule states with preservation of secondary structure content and existent but fluctuating tertiary structure at intermediate-to-slow timescales (µs ms).

## Discussion

TCTP and Mcl-1 have been shown to interact with each other with impacts on their respective cellular stability, yet with opposite outcomes for different cell lines [20, 21, 22]. Zhang et al. [21] observed that Mcl-1 serves as a chaperone to stabilize TCTP in cells, but TCTP has little effect on Mcl-1 cellular levels. In sharp contrast, Liu et al. [20] demonstrated that TCTP stabilizes Mcl-1 in cells by reducing its proteasomal degradation, while Mcl-1 had little effect on TCTP. Giri et al. [20, 21, 22] did similar observation in different cell lines in the context of parasite infection, and further showed that the role of Mule in the TCTP-mediated stabilization of Mcl-1. Here we aimed at characterizing their molecular interactions at an atomic level to further gain knowledge.

We demonstrated that full length TCTP binds to the BH3-binding groove of Mcl-1 and that the interacting region is restricted to the BH3-like motif in fulllength TCTP. We have shown that the TCTP BH3-like motif interchanges between at least two binding modes on a µs-ms timescale in the binding groove of Mcl-1. This particular dynamics can be completely abrogated by a single mutation D16I in the BH3-like motif, with an important increase in affinity to Mcl-1. The D16I mutation at position h1 was introduced to generate a more canonical BH3 sequence. As shown previously [19], a hybrid Bax/TCTP peptide, where the h1 subregion of TCTP was replaced by Bax sequence, totally lost the positive regulation of Bcl-xL anti-apoptotic properties by TCTP and could not rescue haploinsufficient phenotype TCTP+/-thymocytes after γ-irradiation. Both experiments pointed to h1 subregion position of TCTP as central to control TCTP apoptotic function. Here, the rather low affinity of TCTP BH3 for Mcl-1 and the dynamic interface may play a role in Mcl-1-related TCTP biological functions. Among the 51 BH3-only containing proteins reported in BCL2DB [26] (Bcl-2 homologs excluded), TCTP is one of the three proteins having anti-apoptotic properties and is therefore an exception. Further functional studies will allow evaluating the role of D16 in TCTP anti-apoptotic activity and notably if it specifically endows TCTP with an anti-apoptotic role. The R21 residue also lies in the BH3 region of TCTP. The TCTP R21A mutant lost its ability to interact with Mcl-1 in cells [21] and pulse chase experiments further highlighted the faster decay of TCTP R21A compared to the wild type, pointing to the TCTP-Mcl-1 interaction as crucial to maintain high TCTP levels [21]. Our structural model shows electrostatic interactions between the R21 side chain and the Mcl-1 D256 residue in loop α_3_α_4_. This strongly suggests that the loss of the intermolecular R21-D256 salt bridge is sufficient to disrupt the complex. To this regard, it can be hypothesized that the R21 is also crucial for TCTP/Bcl-xL interaction, since TCTP R21A mutant lost the ability to potentiate Bcl-xL [19]. In the Bcl-xL complex, the R21 side-chain interacts with E129 and D133 in Bcl-xL, at similar positions as D256 in Mcl-1. In conclusion, R21, which is strictly conserved in eukaryotic TCTP sequences, but not in prokaryotic sequences[1], appears to play a central role in TCTP interactions with Bcl-2 proteins. On the Mcl-1 side, mutation of K276 to a valine abrogated TCTP/Mcl-1 complex formation [20], but preserved the interaction of Mcl-1 with Bim and Bax. K276 is present at the C-terminus of helix α_4_, on the opposite side of helix α_2_ with respect to the BH3 binding groove, and the K276 side chain creates a salt bridge with D218 in Mcl-1 structure [25]. K276 backbone NMR chemical shifts were not affected upon TCTP binding, confirming that it does not contribute to the interface with TCTP. It is possible that disruption of the K276-D218 salt bridge upon K276V mutation displaces helix α_2_ at the BH3 binding groove and perturbs Mcl-1/TCTP contacts.

Mcl-1 can be rapidly degraded by the proteasome and its rapid turnover is under the control of E3 ligases such as Mule, SCFβ-TrCP, SCFFbw7 and Trim17 [24]. Mule appears to be the major E3 ligase responsible for constitutive degradation of Mcl-1 [66, 62]. The BH3 region of Mule is one of the two identified docking sites of Mule on Mcl-1 for degradation [66, 62, 67]. It is reported that BH3-only peptide (Bak, Bim, Bid, Puma and Noxa) can compete with Mule BH3 at the BH3 binding groove [67]. In addition, a BH3 peptide derived from Bim can prevent Mule to interact with Mcl-1, leading to accumulation of Mcl-1 in cells [66]. Finally, the Mule-Mcl-1 cellular interaction is directly correlated to TCTP levels upon parasite infection [22]. Here we evaluated if TCTP could displace Mule at the Mcl-1 binding groove, as a potential mechanism for the observed TCTP-based protection of Mcl-1 from ubiquitination. Our competition experiments clearly showed that the BH3 peptide of TCTP had much weaker affinity than Mule BH3 peptide for Mcl-1. Considering that full-length TCTP has even lower affinity, this suggests that TCTP protein is not likely to compete with Mule for Mcl-1 at the BH3 binding groove in cells, unless the large TCTP concentration in tumor cells compensates for the very weak binding affinity. This could be the case since TCTP is one of the 20 most abundant protein in eukaryotes [68] with concentration in cancer cells greater than Mule over several orders of magnitude.

The formation of the TCTP/Mcl-1 complex occurs by the docking of the BH3 region on Mcl-1, which results in the destabilization of the TCTP core region into a so-called molten globule. Considering the similarity of TCTP binding to Bcl-xL, this destabilization of TCTP is also highly probable in the TCTP/Bcl-xL complex. Susini et al. [65] suggested that full length TCTP could insert into the mitochondrial membrane upon binding to the Bcl-xL partner using α-helices H2-H3 which have structural homology with Bax α-helices H5-H6. On the other hand, molten-globule intermediates were sometimes reported to enable protein insertion into membranes [69, 70, 71]. We can then imagine than Bcl-2 proteins may facilitate TCTP membrane insertion by destabilizing the TCTP core structure. As exemplified here, the molten globule is more prone to proteolysis in vitro. To some regards, molten globule states can also be considered as a partially disordered or unfolded/misfolded state. In cells, such unfolded states are often recognized by chaperones for later refolding or for proteasomal degradation. The interaction of Mcl-1 with TCTP may then impact the cellular fate of full-length TCTP, through proteasomal degradation or proteolysis, unless specific cellular mechanism protects TCTP. Interestingly, the small chaperon HSP27 stabilizes TCTP levels in prostate cancer cells by preventing its proteasomal degradation. Such ATP-independent small chaperons are known to recognize misfolded proteins. Future investigations will evaluate if HSP27 can specifically recognize, and stabilize, the partially disordered molten globular state of TCTP, triggered for example by Mcl-1 or Bcl-xL binding. Zhang et al. [21] in contrast observed the cellular stabilization of TCTP by Mcl-1, suggesting that the existence of cellular mechanisms compensating for the formation of molten gobule.

We showed that the TCTP_BH3_ peptide has stronger affinity than full-length TCTP for Mcl-1. An in vitro GST pull-down assay showed that a TCTP construct missing the first 10 residues strongly binds Mcl-1 [20]. The first ten amino acids of TCTP fold into the β_1_ strand that is sandwiched between strands β_2_ and β_11_ to form one side of the β-tent [1] in the TCTP structure. Removing the residues 1-10 should therefore largely perturb the overall architecture of TCTP, leading to exposure of the BH3 region. These two in vitro studies are then consistent with an increased binding of TCTP when the BH3 region is readily available for Bcl-2 proteins. This opens the question of potential conformational switch mechanisms in TCTP in the cellular environment prior to Bcl-2 binding. TCTP can be easily phosphorylated by the Plk-1 kinase [72]. We recently showed that TCTP does not show major conformational change upon phosphorylation at position S46 by Plk-1 [32], excluding S46 phosphorylation as a trigger. Bid is a well-known BH3-only protein that is activated by the caspase-8 protease [73, 74]. In Bid, the BH3 region adopts an α-helical structure that becomes available for interaction with Bax and Bcl-2 in its truncated form t-Bid after caspase-8 mediated proteolytic cleavage [73, 74]. TCTP being another BH3-only protein involved in apoptosis, its cleavage by caspase could be considered. TCTP activation through peptidic cleavage has already been discussed in other contexts [75, 76]. Indeed, TCTP interacts with the caspase recruitment domain of the Apaf-1 apoptosome to inhibit the amplification of caspase cascade [76]. However only a C-terminally cleaved TCTP is able to bind Apaf-1. The TCTP sequence does not contain any known caspase enzymatic site, and protease inhibitors could not help identifying the involved protease activity for TCTP activation [76]. Another study pointed out that recombinant caspase-3 could not cleave TCTP [75]. Taken together, whether, and how, TCTP might be activated for Bcl-2 binding by post-translational modification (phosphorylation, cleavage, binding to other partners) remains to be investigated. The burying of TCTP BH3-like motif in its globular domain might indeed represent an elegant strategy to preserve normal cell functioning, while the deregulation of the apoptosis pathway associated to tumorigenesis may drive the activation of TCTP anti-apoptotic effect by exposing its BH3 to partners.

TCTP has become an attractive drug target in cancer and other diseases. TCTP functions by interaction with numerous other proteins [1] and blocking the TCTP interactome might be a viable therapeutical strategy. This is exemplified by the sertraline drug that was proposed to prevent TCTP from binding MDM2, in order to restore high p53 levels. Small compounds targeting TCTP/Mcl-1 or TCTP/Bcl-xL complexes may also be potent alternatives to reduce TCTP pro-survival properties. TCTP is a globular protein with no obvious binding pockets, questioning its druggability by conventional approaches. Yet, in silico drug screening led to promising compounds [77, 78, 79], although the binding mode remain to be better described for further improvement. Our present work shed the light on biophysical properties of TCTP when bound to Mcl-1. Full-length TCTP slowly binds Mcl-1, while the BH3 peptide rapidly binds. This indicates a rare rate-limiting conformational switch event allowing the population of a binding competent state. Obviously, small compounds preventing or facilitating this event would have the potential to block or speed up complex formation, hence controlling TCTP Bcl-2-mediated apoptotic effects. We are currently investigating this conformational switch. Alternatively, compounds (de)stabilizing TCTP into a molten globule-state could play a role in the cellular stability of the protein and might represent an alternative to antisense oligonucleotide strategy [14] to reduce TCTP levels in tumor cells.

As a conclusion, TCTP is a small but highly versatile protein that can undergo a major structural and dynamic reorganization upon Mcl-1 binding. Our work improved knowledge of its plasticity to adapt to various reported partners [1] in the context of TCTP biology and potential future drug design strategies.

This article contains supporting information.

## Acknowledgements

We thank Nadine Assrir for sharing her protocol and knowledge for TCTP preparation.

## Funding and additional information

The FP7 WeNMR (project #261572), H2020 West-Life (project #675858), the EOSC-hub (project #777536) and the EGI-ACE (project #101017567). European e-Infrastructure projects are acknowledged for the use of their web portals, which make use of the EGI infrastructure with the dedicated support of CESNET-MCC, INFN-PADOVA-STACK, INFN-LNL-2, NCG-INGRID-PT, TW-NCHC, CESGA, IFCA-LCG2, UA-BITP, SURFsara and NIKHEF, and the additional support of the national GRID Initiatives of Belgium, France, Italy, Germany, the Netherlands, Poland, Portugal, Spain, UK, Taiwan and the US Open Science Grid. Financial support from the IR INFRANALYTICS FR2054 for conducting the research is gratefully acknowledged.This work was supported by the French Infrastructure for Integrated Structural Biology (FRISBI) ANR-10-INBS-0005. We also thank the synchrotron facility SOLEIL (St. Aubin) for allocating beam time (proposal 20181072) and its dedicated staffs for technical help with the beamline SWING. This work is supported by the “IDI 2016” project funded by the IDEX Paris-Saclay ANR-11-IDEX-0003-02 as a PhD fellowship to F.M.

