## Supplementary Information for "Structural transitions in TCTP tumor protein upon Mcl-1 binding"

### Supporting Information for:
### Structural transitions in TCTP tumor protein upon Mcl-1 binding

#### Supplementary figures

```
TCTP Full Length (TCTP FL)
-1  1
 GP MIIYRDLISH DEMFSDIYKI REIADGLCLE VEGKMVSRTE GNIDDSLIGG NASAEGPEGE GTESTVITGV

    DIVMNHHLQE TSFTKEAYKK YIKDYMKSIK GKLEEQRPER VKPFMTGAAE QIKHILANFK NYQFFIGENM
                                     172
    NPDGMVALLD YREDGVTPYM IFFKDGLEME KC

Mcl-1 ΔPESTΔTM
-1  1 (172)
 GP DELYRQSLEI ISRYLREQAT GAKDTKPMGR SGATSRKALE TLRRVGDGVQ RNHETAFQGM LRKLDIKNED

    DVKSLSRVMI HVFSDGVTNW GRIVTLISFG AFVAKHLKTI NQESCIEPLA ESITDVLVRT KRDWLVKQRG
                    156 (327)
    WDGFVEFFHV EDLEGG
```

Figure S1: **Primary sequence of FL-TCTP and Mcl-1 ΔPEST ΔTM used for interaction studies.** For TCTP Full Length (FL-TCTP), the N-Terminal GP residues (grey) originate from the TEV cleavage site. Residues M1 to C172 corresponds to the full length sequence of human TCTP protein (UniProtKB P13693). For Mcl-1 ΔPEST ΔTM, the N-Terminal GP residues originate from the Prescission cleavage site. Residues D1 to G156 corresponds to the D172 to G327 segment of human Mcl-1 (UniProtKB Q07820).

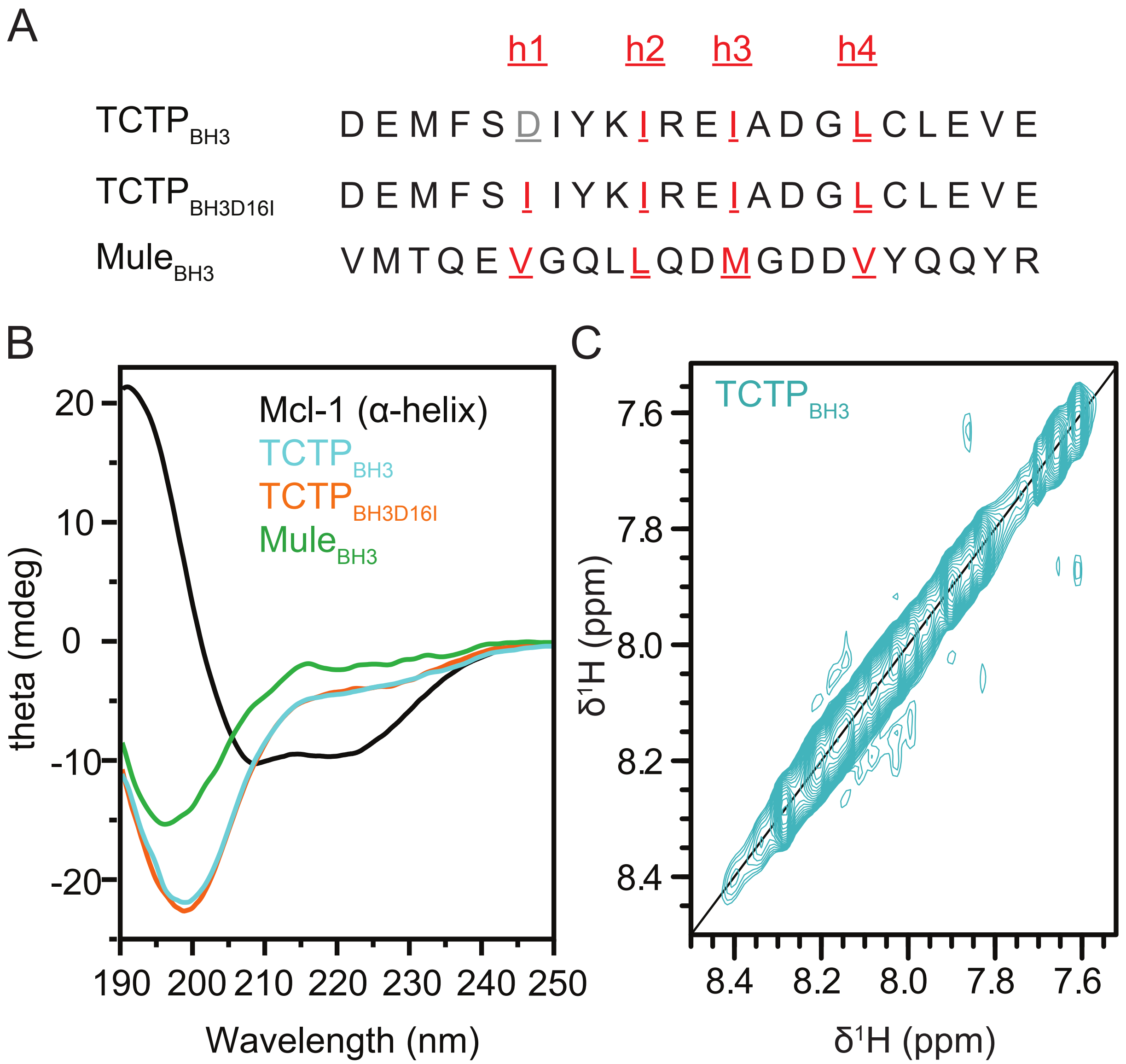


Figure S2: **Primary sequences and structural characterization of BH3 peptides related to TCTP and Mule.** (A) Primary sequence of the TCTP BH3-like peptide ($TCTP_{BH3}$), the canonical D16I mutant ($TCTP_{BH3D16I}$) and the canonical BH3 peptide from the E3 ubiquitine ligase Mule ($Mule_{BH3}$). Conserved position h1, h2, h3, h4 in canonical BH3 motifs are highlighted (red). $TCTP_{BH3}$ and $Mule_{BH3}$ peptide constructs correspond to the segment 11-32 and 1971-1992 in the full-length TCTP and Mule, respectively. (B) Far-UV (190-250 nm) CD experiments with $TCTP_{BH3}$ (100 µM, cyan), $TCTP_{BH3D16I}$ (100 µM, orange), $Mule_{BH3}$ (100 µM, green) and an α-helical reference Mcl-1 ΔPEST ΔTM (Mcl-1) (100 µM, black). Experiments were carried out at 298 K in 2.5 mM phosphate buffer pH 6.5. (C) $^1H$-$^1H$ NOESY spectrum ($\tau_m$ = 200 ms) of $TCTP_{BH3}$. Experiment was recorded at 800 MHz and 278 K in the following buffer: 50 mM MES pH 6.5, 50 mM NaCl, 2 mM TCEP in 5 % $D_2O$ / 95 % $H_2O$.

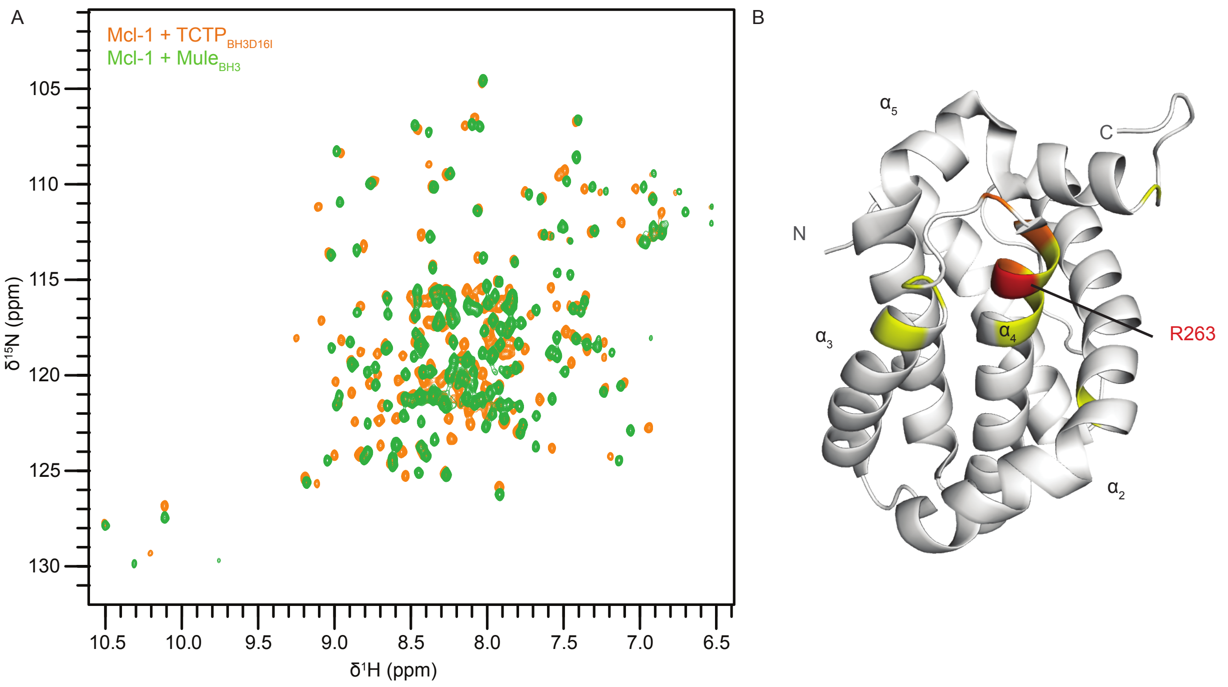


Figure S3: **Comparison of $\mathbf{TCTP_{BH3D16I}}$ and $\mathbf{Mule_{BH3}}$ binding to Mcl-1.** (A) Overlay of $^{15}$N SOFAST HMQC spectra from $^{15}$N-Mcl-1 ΔPEST ΔTM (Mcl-1) (100 µM) in complex with TCTP BH3-like peptide D16I mutant ($TCTP_{BH3D16I}$) (orange) or $Mule_{BH3}$ (green). (B) Mapping of combined $^{1}$H-$^{15}$N chemical shift perturbations between $^{15}$N-Mcl-1 ΔPEST ΔTM (Mcl-1) (100 µM) in complex with TCTP BH3-like peptide D16I mutant ($TCTP_{BH3D16I}$) or $Mule_{BH3}$ ($\Delta\delta_{comb} > 0.9$, red; $0.9 > \Delta\delta_{comb} > 0.65$, orange; $0.65 > \Delta\delta_{comb} > 0.4$, yellow; $0.4 > \Delta\delta_{comb}$, white) on the NMR structure of Mcl-1 [80]. Experiments were recorded at 950 MHz and 308 K in the following buffer: 50 mM EPPS pH 8, 50 mM NaCl, 2 mM TCEP in 5 % $D_2O$ / 95 % $H_2O$.

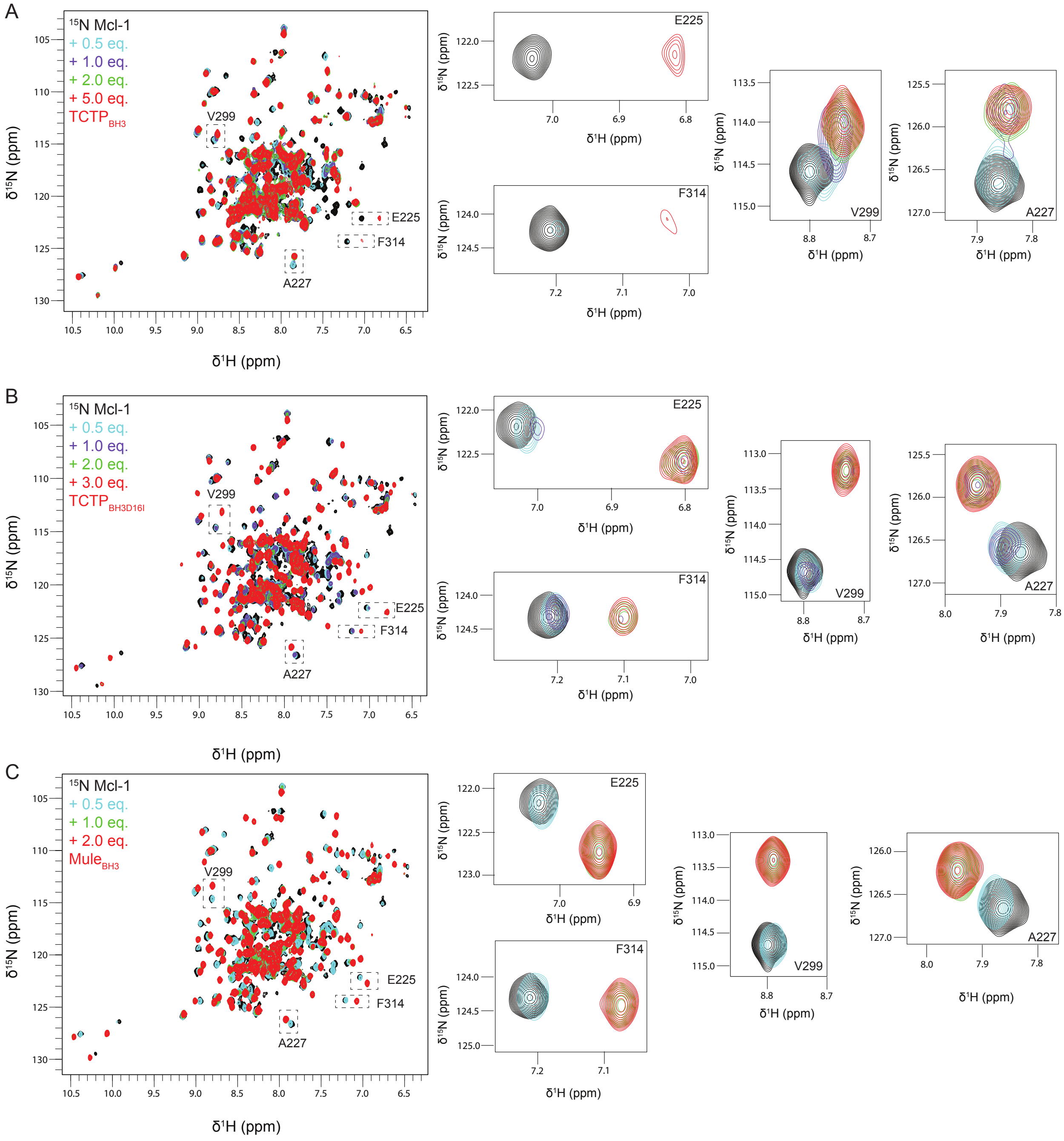


Figure S4: **NMR titrations of Mcl-1 with BH3-derived peptides and exchange regimes.** Overlay of $^{15}$N SOFAST HMQC spectra from $^{15}$N-Mcl-1 ΔPEST ΔTM (Mcl-1) (100 µM) and upon addition of increasing amount of (A) TCTP BH3-like peptide (TCTP$_{BH3}$) or (B) TCTP BH3-like peptide D16I mutant (TCTP$_{BH3D16I}$) or (C) Mule BH3 peptide (Mule$_{BH3}$). A close-up view for NMR crosspeaks corresponding to residues E225, F314, V299 and A227 is given to illustrate the exchange regime along the titration. Experiments were recorded at 950 MHz and 298 K in the following buffer: 50 mM MES pH 6.5, 50 mM NaCl, 2 mM TCEP in 5 % $D_2O$ / 95 % $H_2O$.

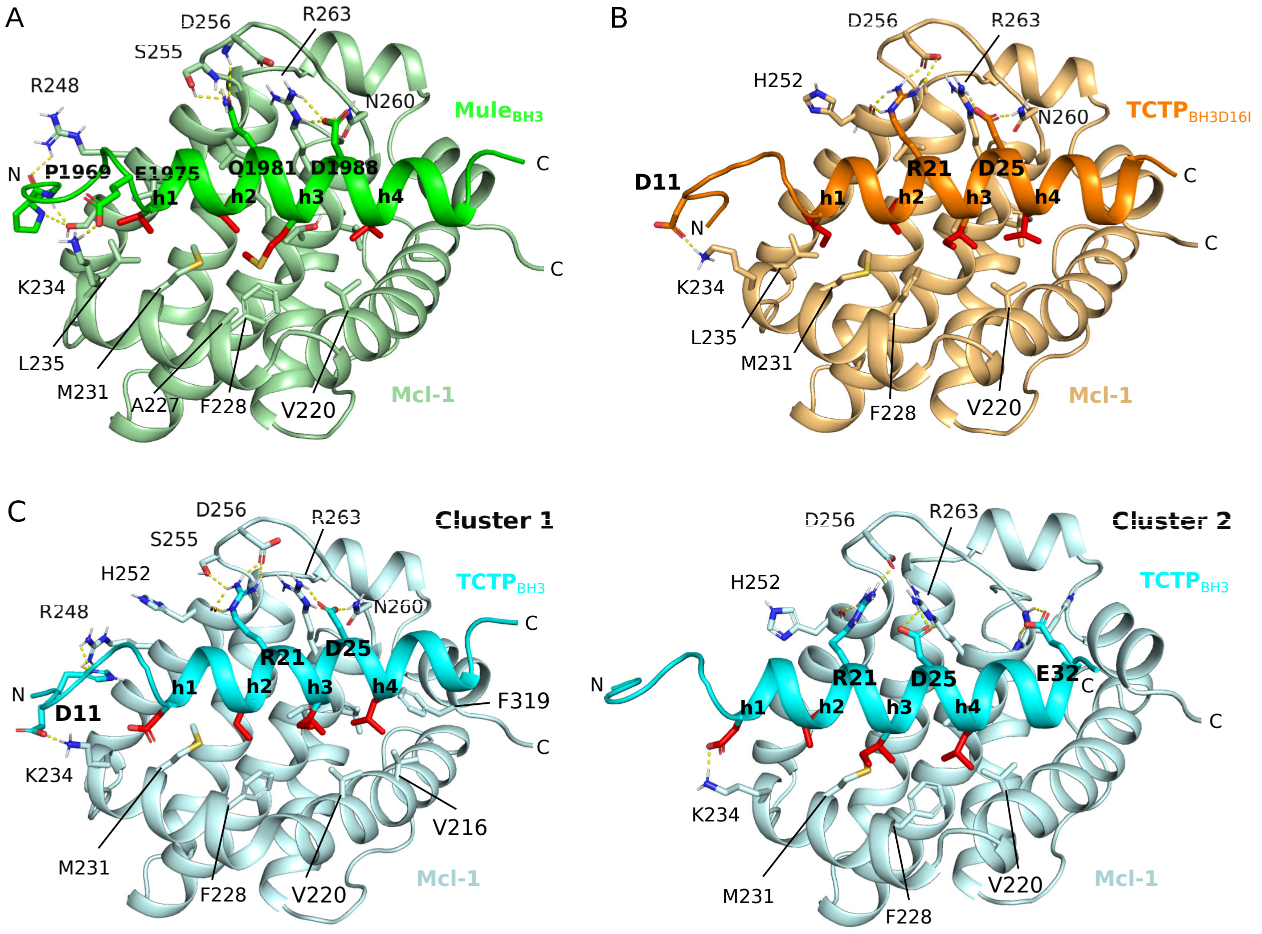


Figure S5: **Interaction interfaces between Mcl-1 and BH3 peptides in HADDOCK docking models.** (A) Representative structure of the top ranked cluster for Mcl-1/Mule$_{\mathrm{BH3}}$. (B) Representative structure of the top ranked cluster for Mcl-1/TCTP$_{\mathrm{BH3D16I}}$. (C) Representative structure of (left) cluster 1 and (right) cluster 2 for Mcl-1/TCTP$_{\mathrm{BH3}}$. In (A), (B) and (C), bold and regular labels indicate BH3 and Mcl-1 residues, respectively. In BH3 peptides, residues in h1-4 positions (red sticks) and surrounding hydrophobic residues in Mcl-1 (sticks) are highlighted. Electrostatic contacts (yellow dashed lines) and contributing residues (sticks) are highlighted.

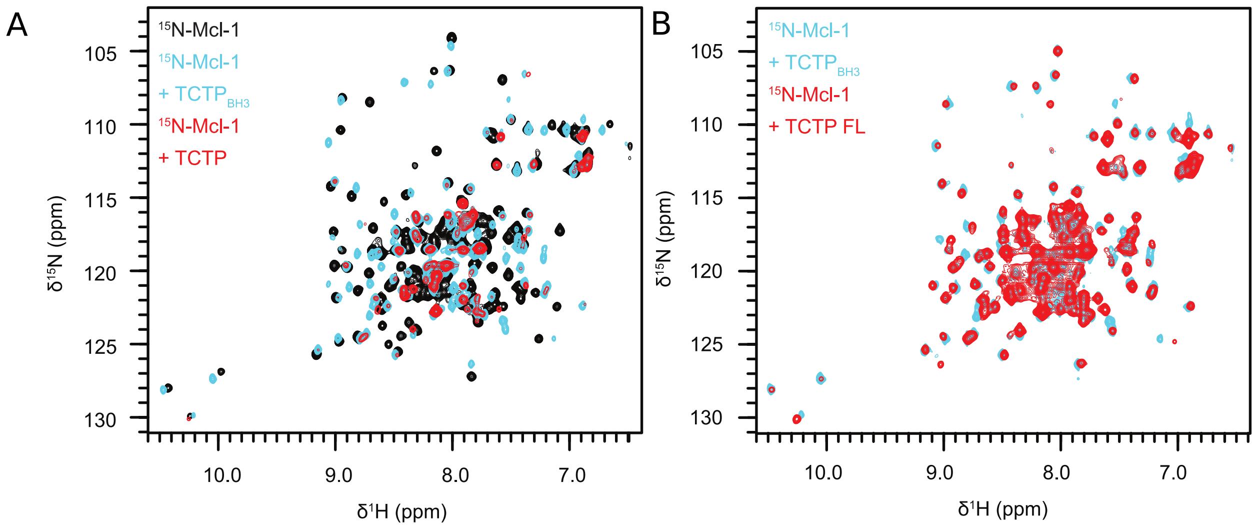


Figure S6: **Comparison of TCTP and $\mathbf{TCTP_{BH3}}$ binding to Mcl-1.** (A) Overlay of $^{15}$N SOFAST HMQC spectra from isolated $^{15}$N-Mcl-1 (100 µM, black) and in complex with TCTP BH3-like peptide ($TCTP_{BH3}$) (cyan) or full length TCTP (red). (B) Overlay of $^{15}$N SOFAST HMQC spectra from $^{15}$N-Mcl-1 ΔPEST ΔTM (Mcl-1) (100 µM) in complex with TCTP BH3-like peptide ($TCTP_{BH3}$) (cyan) or full length TCTP (red). Spectral intensity was scaled for comparison of chemical shift profiles in both conditions. Experiments were recorded at 950 MHz and 308 K in the following buffer: 50 mM EPPS pH 8.0, 50 mM NaCl, 2 mM TCEP in 5 % $D_2O$ / 95 % $H_2O$.

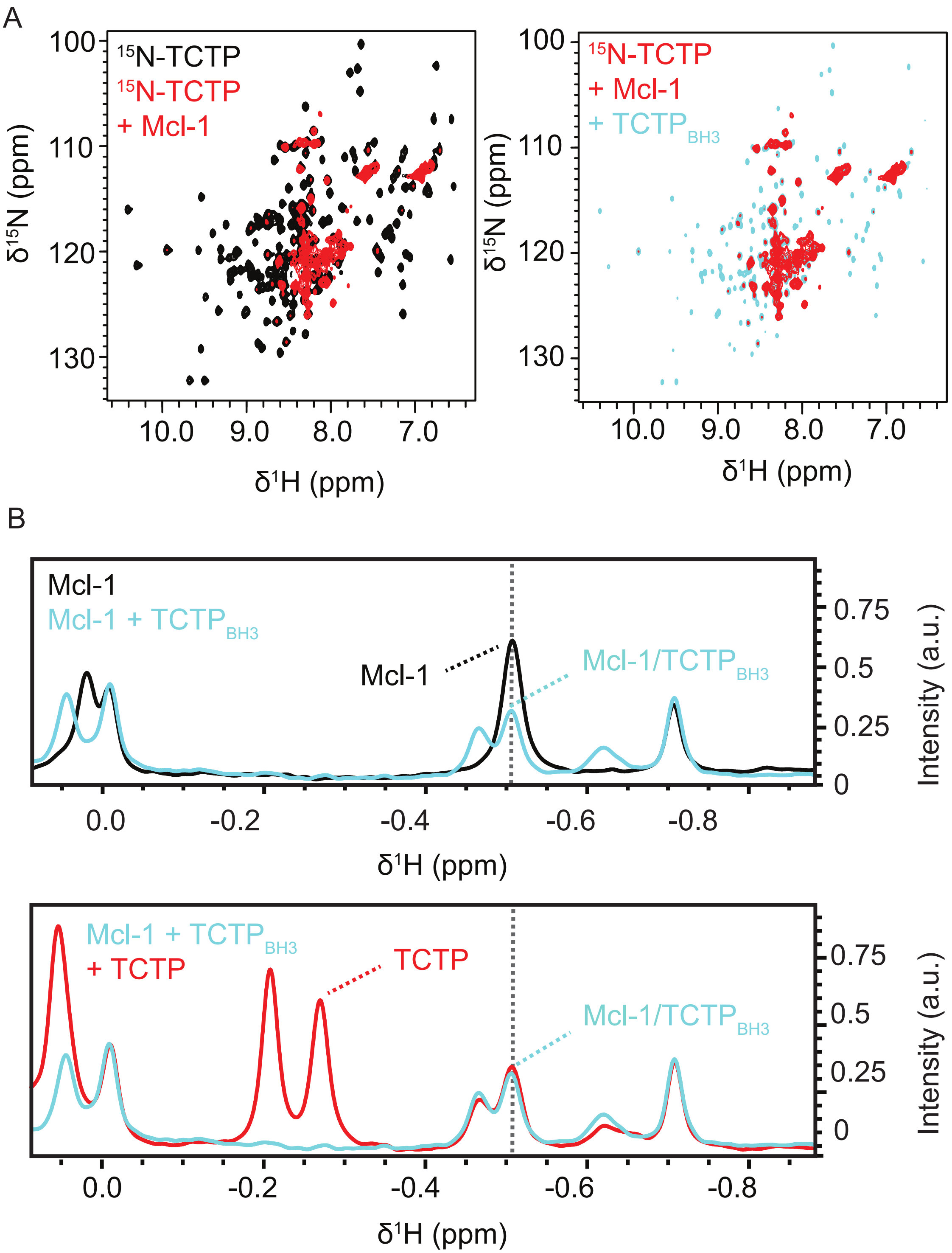


Figure S7: **Relative binding capacity of $\mathbf{TCTP_{BH3}}$ and FL-TCTP to Mcl-1.** (A) Overlay of $^{1}$H spectra from isolated Mcl-1 ΔPEST ΔTM (Mcl-1) (100 µM, black), upon addition of $TCTP_{BH3}$ (4 eq., blue), FL-TCTP (2 eq., red), and after incubation time (2 hrs) (green). (B) Overlay of $^{15}$N SOFAST HMQC spectra from isolated $^{15}$N-TCTP (100 µM, black), in complex with unlabeled Mcl-1 ΔPEST ΔTM (Mcl-1) (2 eq.) (red) and upon addition of $TCTP_{BH3}$ (2 eq.) (cyan). Experiments were recorded at 950 MHz and 308 K in the following buffer: 50 mM EPPS pH 8, 50 mM NaCl, 2 mM TCEP in 5 % $D_2O$ / 95 % $H_2O$.

#### Supplementary tables

| | Mcl-1 | TCTP/Mcl-1 |
|---|---|---|
| Organism | Human | Human |
| Source | E. Coli | E. Coli |
| Description - sequence (including tags) + bound ligands/modifications, etc. | See Supp. Fig. S1 | See Supp. Fig. S1 |
| Extinction coefficient | 19480 $M^{-1}.cm^{-1}$ | 19480 + 11920 = 31400 $M^{-1}.cm^{-1}$ |
| *M* from chemical composition | 17765.21 Da | 17765.21 + 19652.39 = 37417.6 Da |
| For SEC-SAS, loading volume/concentration, flow rate | 75µL/500µM, 0.5mL/min | 75µL/500µM, 0.5mL/min |
| Solvent details | 50 mM CHES pH 9, 50 mM NaCl and 2 mM TCEP | 50 mM CHES pH 9, 50 mM NaCl and 2 mM TCEP |
| Guinier Analysis | Mcl-1 | TCTP/Mcl-1 |
| *I(0)* | 0.0078 +/- 0.00001 | 0.0036 +/- 0.00001 |
| $R_g$ | 1.758 +/- 0.004 nm | 2.102 +/- 0.008 nm |
| $qR_g$ range | 0.14-1.3 | 0.22-1.29 |
| *M* from *I*(0) (ratio to expected value) | 21.7 kDa (1.22) | 34.7 kDa (0.93) |
| *P*(*r*) analysis | Mcl-1 | TCTP/Mcl-1 |
| $V_c$ | 206 | 283 |

Table S1: **Reporting of essential Small Angle Scattering (SAS) data acquisition, sample details, data analysis, modelling fitting and software used.** We used a reporting template from [81] to provide relevant details of experiments performed with biomolecules in solution at SOLEIL synchrotron (Saint-Aubin, France).

#### Supplementary files

##### Supplementary file S1

Download link: https://mycore.core-cloud.net/index.php/s/TqNE29tYp95lljY

Mcl1_BH3s.zip: CcpNmr [44] session with $^{1}$H-$^{15}$N SOFAST HMQC spectra of isolated $^{15}$N-Mcl-1 ΔPESTΔTM (Mcl-1) and in complex with TCTP BH3-like peptide ($TCTP_{BH3}$) or TCTP BH3-like peptide D16I mutant ($TCTP_{BH3D16I}$) or the E3 ubiquitine ligase Mule BH3 peptide ($Mule_{BH3}$). The corresponding backbone assignments are provided within the CcpNmr session. Experiments were recorded at 950 MHz and 308 K in the following buffer: 50 mM MES pH 6.5, 50 mM NaCl, 2 mM TCEP in 5 % $D_2O$ / 95 % $H_2O$.
